# A CRISPRi/a screening platform to study cellular nutrient transport in diverse microenvironments

**DOI:** 10.1101/2023.01.26.525375

**Authors:** Christopher Chidley, Alicia M. Darnell, Benjamin L. Gaudio, Evan C. Lien, Anna M. Barbeau, Matthew G. Vander Heiden, Peter K. Sorger

## Abstract

Blocking the import of nutrients essential for cancer cell proliferation represents a therapeutic opportunity, but it is unclear which transporters to target. Here, we report a CRISPRi/a screening platform to systematically interrogate the contribution of specific nutrient transporters to support cancer cell proliferation in environments ranging from standard culture media to tumor models. We applied this platform to identify the transporters of amino acids in leukemia cells and found that amino acid transport is characterized by high bidirectional flux that is dependent on the composition of the microenvironment. While investigating the role of transporters in cystine starved cells, we uncovered a novel role for serotonin uptake in preventing ferroptosis. Finally, we identified transporters essential for cell proliferation in subcutaneous tumors and found that levels of glucose and amino acids can restrain proliferation in that environment. This study provides a framework for the systematic identification of critical cellular nutrient transporters, characterizing the function of such transporters, and studying how the tumor microenvironment impacts cancer metabolism.

## INTRODUCTION

Altered cellular metabolism is a common feature of tumors, enabling cancer cells to maximize proliferation in nutrient-limited environments^1^. Cellular adaptations to the tumor environment (TME) generally include an increase in the rate of nutrient uptake, a diversification of uptake mechanisms, and a rewiring of intracellular metabolism so tumors can more efficiently convert available nutrients into biomass^2^. Amino acids contribute substantially to the formation of cellular biomass^3^, and many cancers are dependent on environmental supplies of non-essential amino acids for growth and survival. For example, acute lymphoblastic leukemias are dependent on asparagine, luminal breast cancers and breast cancer brain metastases are auxotrophic for serine^2,4–7^, and yet other cancers silence arginine biosynthesis or upregulate glutamine metabolism. Such dependencies can potentially be targeted therapeutically by inhibiting nutrient uptake^8^.

Nutrients are actively transported across cell membranes by a large class of transporter proteins called solute carriers (SLCs)^9^. The activity of many SLCs is influenced by the nutrient composition of the microenvironment^10^ and studying the function of these SLCs, or determining which ones might be relevant therapeutic targets, requires characterizing transport under physiologically relevant conditions. Unfortunately, much of our current knowledge of SLCs derives from experiments in cell-free systems and many SLC substrates identified in these experiments (hereafter “annotated” substrates) have not been evaluated in cells or in vivo^11^. For example, over 60 SLCs are annotated as being amino acid transporters but it is not clear which are the dominant or growth-limiting transporters used by cells under specific conditions^11–13^. Recent studies have also uncovered the transporters for essential metabolites such as NAD^+14–16^ and glutathione^17^, but 20–30% of SLCs, including many that are essential in human cells, lack a known substrate or metabolic function^9,18^. The development of CRISPR screening^19^ and efforts to better characterize the composition of the TME^20,21^ present opportunities to better define the role of nutrient transporters in human cells.

In this paper we describe the use of CRISPR interference (CRISPRi) and activation (CRISPRa) screening using custom sgRNA libraries to systematically interrogate the individual contributions to nutrient uptake or release of all SLC and ATP-binding cassette (ABC) transporters found in human cells. We determined the transporters responsible for amino acid import in K562 leukemia cells and used mass spectrometry-based transport assays to reveal that amino acid transport is characterized by high bidirectional flux at the plasma membrane and is highly sensitive to environmental conditions. We also used CRISPRi to identify transporters essential for cell proliferation in standard and physiological growth media, and in subcutaneous tumors in mice, and identified nutrients that underlie condition-specific essentialities to infer transporter function. Finally, we used CRISPRa to identify nutrients limiting for cellular proliferation in different environments, providing a framework for characterizing relevant cell nutrient transporters and how their interaction with the tumor microenvironment shapes cancer metabolism.

## RESULTS

### Cell-based screens for amino acid transporters

To assess transporter function in cells, we used a dual approach: transporter knockdown (KD) via CRISPRi and transporter overexpression (OE) via CRISPRa. Based on previous data showing that transporter genes are often among the top hits in CRISPR screens^19,22–24^, we hypothesized that KD/OE of transporters would change the rate of substrate transport and that changes could be scored using growth-based pooled screens (**Figure 1A**). More specifically, we postulated that when cells are cultured in conditions in which a nutrient is limiting for proliferation, KD of a transporter gene needed for import of that nutrient would reduce proliferation and, conversely, OE of a gene whose product was capable of importing that nutrient would increase proliferation. Using an analogous logic, transporters participating in nutrient export will display opposite phenotypes to importers in such screens. We engineered chronic myeloid leukemia (K562) cells to express either dCas9-KRAB (for CRISPRi) or dCas9-SunTag (for CRISPRa). To validate CRISPR-based transporter KD or OE, we targeted three SLC7 amino acid transporters expressed in K562 cells (SLC7A1, SLC7A5, SLC7A6) and three that were not expressed (SLC7A2, SLC7A3, SLC7A7). We found that CRISPRi/a of transporters resulted in decreases or increases in mRNA expression that were strong and specific (**Figure 1B**).

**Figure 1.**
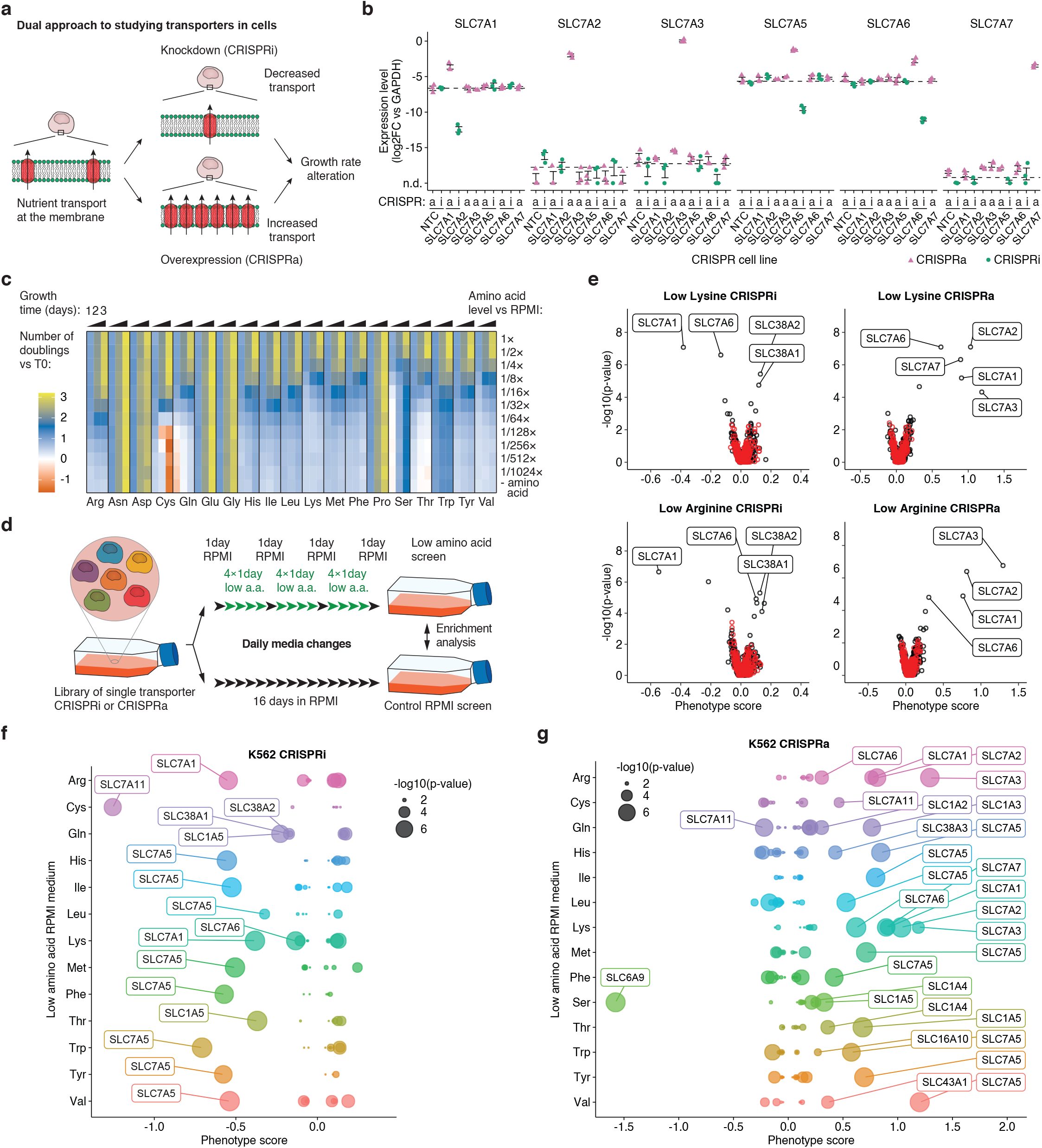
CRISPRi/a screens identify the transporters of amino acids in K562 cells. (a) Cartoon of the general approach used to identify transporters in cells. Individual transporter genes are knocked down via CRISPRi or overexpressed via CRISPRa, and modified transport activity is detected by changes in proliferation. (b) CRISPRi/a of genes in the SLC7 amino acid transporter family leads to specific perturbation of gene expression. Gene expression in K562 CRISPRi/a cells with specific or non-targeting control (NTC) sgRNAs was quantified by RT-qPCR relative to the housekeeping gene GAPDH. Error bars: ± SEM of n = 3 technical replicates. n.d., not detected. (c) Identification of amino acids that limit proliferation of K562 cells when their medium level is reduced. Data were determined using a luminescent cell viability assay and represent the average log2FC in luminescence relative to time = 0 days (T0). (d) Cartoon of pooled screening strategy used to identify amino acid transporters. Library pools were grown in RPMI medium where a specific amino acid was present at a level that reduced proliferation by 50% relative to complete RPMI (low a.a.). (e) Volcano plots of transporter CRISPRi/a screens in K562 cells in low lysine and low arginine. Phenotype scores represent averaged and normalized sgRNA enrichments in low a.a. versus RPMI and -log10(p-value) were determined using a Mann-Whitney test of sgRNA enrichments compared to all NTC sgRNAs. Black circles represent transporter genes and red circles represent negative control genes generated by randomly grouping sets of NTC sgRNAs. Data were computed from two screen replicates. (f) Bubble plots displaying CRISPRi screen scores determined for all 64 transporters annotated as capable of amino acid transport^12^. (g) same as (f) for CRISPRa screens.

Next we constructed CRISPRi and CRISPRa sgRNA pooled lentiviral libraries targeting all 489 annotated members of the SLC and ABC transporter families (hereafter referred to as “transporters”)^18,25,26^; the library comprised 10 sgRNAs per gene and 730 non-targeting control (NTC) sgRNAs (Table S1 and Methods; plasmid pools will be made available). The primary known function of ABC transporters is to export xenobiotics from cells, but some export metabolites, such as glutathione^27^ and nucleoside analogs^28^, suggesting that ABC transporters also contribute to nutrient homeostasis. Libraries of single transporter KD or OE cells were subsequently prepared by lentiviral transduction of K562 CRISPRi or CRISPRa parental lines (**Figure S1A**).

To identify conditions that would enable selection for transporter phenotypes, we identified all amino acids in RPMI-1640 (RPMI) culture medium whose absence limited K562 cell proliferation. This was done by performing proliferation assays with a series of single amino acid dropout RPMI media to which amino acids were added back to between 0.1% and 100% of their normal levels (**Figure 1C**). For 13 of the 19 amino acids found in RPMI, a reduction in media levels decreased proliferation, consistent with previous evidence demonstrating net consumption of these amino acids by mammalian cells^3^. Removing the amino acids Asn, Asp, Glu, Gly, and Pro from the medium did not affect proliferation, so we were unable to identify transporters for these non-essential amino acids. K562 cells exhibited an early response to low Ser but proliferation rates subsequently returned to normal, most likely as a consequence of increased expression of serine biosynthesis enzymes^29^. We were therefore only able to screen for serine transporters under transient and mild growth limitation conditions. Cystine deprivation was the only condition we identified that induced significant cell death (**Figure 1C**), likely through ferroptosis^30,31^.

We performed pooled CRISPRi/a screens at concentrations of amino acid that reduced K562 cell proliferation by ~50% for all 13 growth-limiting amino acids (**Table S2,3**). This made it possible to identify transporter perturbations that caused either a decrease or increase in proliferation. Libraries of single transporter KD or OE cells were subjected to three successive rounds of low amino acid exposure lasting for four days each with daily media changes followed by one recovery day in complete medium (**Figure 1D**). Perturbations causing changes in proliferation in low amino acid conditions were identified by comparison to a paired control screen performed in complete RPMI medium (**Figure 1D**). Phenotypes were quantified using two gene-level scores^32^: a phenotype score representing the average sgRNA enrichment/depletion, and a confidence score representing the estimated significance of the enrichment/depletion as compared to NTC sgRNAs (Methods)^23^.

Amino acid limitation can activate the GCN2 starvation response and repress mTOR activity, and this response can result in changes in transporter gene expression^12^. To assess the extent to which this occurred under screening conditions we used Western blotting to assay phosphorylation of p70 S6 kinase and 4E-BP1 by mTORC1, and eIF2α by GCN2 (**Figure S1B**). We found that, when K562 cells were cultured in low Arg, Lys, or His screening conditions, the GCN2 response was activated to a degree that was intermediate between the basal activity observed in cells grown in complete medium and the full response elicited by complete starvation for these amino acids. Refreshing the medium after one day relieved GCN2 activation in low amino acid conditions but not in complete starvation. A similar pattern of activity was observed for mTOR: partial mTOR repression in screening conditions with reactivation following media exchange only for cells in low amino acid conditions (**Figure S1B**). We conclude that screening conditions have the potential to activate the GCN2/mTOR pathways and thus alter transporter gene expression, and that components of these pathways might be expected to score as hits. However, we hypothesized that this would not substantially interfere with our identification of additional and likely stronger phenotypes arising from KD/OE of specific transporter proteins.

### CRISPRi/a screening results

We found that CRISPRi screens in low amino acid conditions were enriched in hits with negative phenotype scores, consistent with a role for these transporters in net amino acid import. For all 13 amino acid-limited screening conditions we identified at least one significantly depleted transporter gene (**Figure 1E,F** and **Figure S1C**). Screens with one strong negative hit are consistent with the idea that a single transporter is responsible for the bulk of the import of the limiting amino acid. For example, we found that SLC7A5 (also known as LAT1)^33^ is likely the primary importer for all large neutral amino acids (His, Ile, Leu, Met, Phe, Trp, Tyr, and Val) when they are limiting for growth^12,26^. Screens with more than one negative hit are consistent with a role for multiple partially redundant transporters; this was observed in the case of CRISPRi of SLC1A5, SLC38A1, and SLC38A2 in glutamine-limiting conditions (**Figure 1F**).

CRISPRa screens in low amino acid conditions primarily yielded hits with positive scores, consistent with an increase in amino acid import (**Figure 1G** and **Figure S1C**). CRISPRa identified more hits than CRISPRi since it is applicable to all transporters in the genome, not only the ~50%^34^ that are well expressed in parental K562 cells. For example, SLC7A1, SLC7A2 and SLC7A3 (also known as CAT1, CAT2, and CAT3) are annotated as transporters of arginine and lysine. They were strong hits in both low Lys and low Arg CRISPRa screens, but among these transporters, only SLC7A1 was a hit in the CRISPRi screen, reflecting the fact that only SLC7A1 is detectably expressed in K562 cells (**Figure 1B,E**). Thus, SLC7A1 is the main transporter for Lys and Arg in K562 cells, but SLC7A2 and SLC7A3 are capable of transporting these amino acids if expressed.

Scores in CRISPRi/a screens were highly comparable across biological replicates, suggesting that our screening approach was reproducible (**Figure S1D–F**). We found that all strong CRISPRi hits were also hits in the corresponding CRISPRa screens, confirming that transport was not saturated under screening conditions at endogenous levels of gene expression. We also found that many transporter OE and KD were associated with insignificant phenotypic scores. Insignificant scores could have either technical or biological explanations. An example of a technical explanation would be absence of a measurable phenotype because sgRNA introduction did not appreciably affect expression of a cognate transporter. However, in those cases in which a significant phenotype was reproducibly observed for a gene in at least one screening condition (demonstrating that KD or OE for that gene was effective), it was possible to interpret negative phenotypes under other conditions. For example, CRISPRa screens showed that SLC1A2 can transport glutamine and the presence of this positive result means that negative findings under other CRISPRa screening conditions can be interpreted as evidence of SLC1A2 inactivity against other amino acids tested (**Figure S1C**).

To more thoroughly explore the effects of transporter perturbation across multiple conditions, we constructed individual cell lines from K562 CRISPRi/a parental lines each of which expressed a single sgRNA identified in screens; transporter KD/OE was confirmed by RT-qPCR. We then measured growth phenotypes in competition assays in which an sgRNA-expressing KD/OE cell line was mixed in roughly equal proportion with a control cell line expressing an NTC sgRNA. The mixture of cells was cultured for 10-14 days and the fraction of each cell line in the culture was then determined by qPCR (**Figure S2A,B**). We first used this approach to validate results from primary screens using seven genes in the SLC7 family having positive, negative or no phenotypes in low His, Lys, and Arg conditions; we obtained results that were highly concordant with those obtained in pooled library screens (**Figure 2A**). For example, we observed that OE of SLC7A7, which is annotated as a transporter of Arg and Lys^33^, resulted in a proliferation advantage in low Lys but not in low Arg in both screens and competition assays (**Figure 2A**). Because SLC function is influenced by the intra- and extracellular substrate concentrations, we asked whether the absence of growth phenotype for SLC7A7 OE in low Arg (8 μM) was due to competition for Lys (220 μM) in the medium (**Table S2**). However, when we measured growth scores in RPMI in which the levels of Lys had been reduced 4-fold, we observed no changes in phenotype. These data suggest that SLC7A7 imports Lys but not Arg under growth-limiting concentrations (**Figure 2A**).

**Figure 2.**
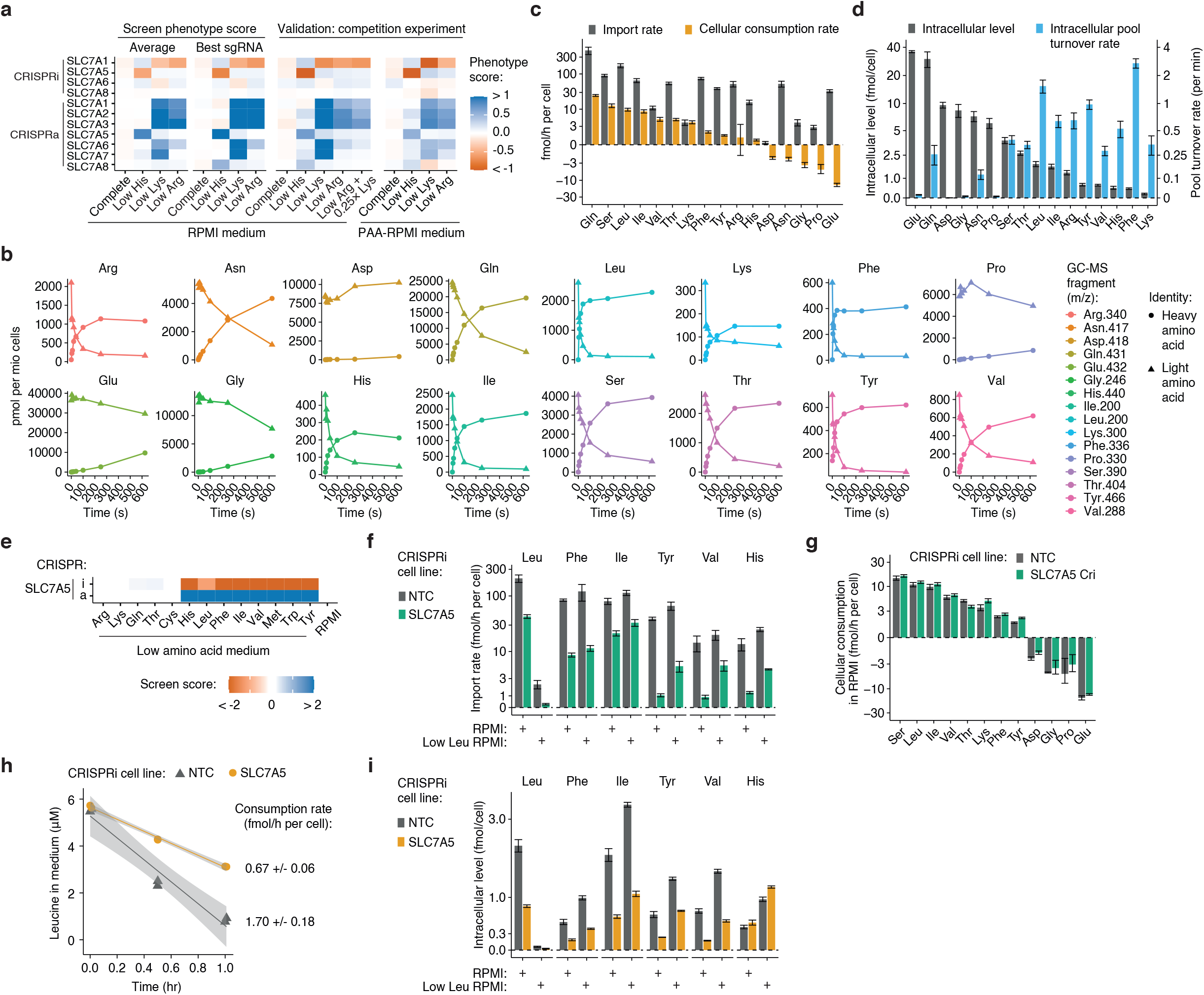
Measurement of amino acid transport rates reveals that CRISPRi of SLC7A5 specifically reduces transport of large neutral amino acids. (a) Phenotype scores obtained in transporter CRISPRi/a screens for SLC7 family genes were validated in competition assays in RPMI and in RPMI with amino acids adjusted to human plasma levels (PAA–RPMI). K562 CRISPRi/a expressing sgRNAs with the strongest phenotype score determined in screens (best sgRNA) were mixed 1:1 with cells expressing NTC sgRNAs, and phenotype scores were calculated from qPCR determination of the ratio after 10 days in complete or in low amino acid medium. (b) Time course of amino acid import into K562 cells determined by following the intracellular accumulation of stable heavy-isotope labeled amino acids by GC-MS. (c) Amino acid import and cellular consumption rates of K562 cells growing in RPMI. Import rates were determined by linear regression of the early phase of heavy-isotope labeled amino acid accumulation. Consumption rates were determined by linear regression over time of amino acid levels in the growth medium of K562 cells. Data are mean ± SEM of n = 6 experiments for import and n = 5 for consumption. (d) Intracellular amino acid levels (n = 7 ± SEM) and pool turnover rates of K562 cells growing in RPMI. Pool turnover rates were inferred by dividing amino acid import rates by intracellular levels. (e) Screen scores for K562 SLC7A5 CRISPRi/a. (f) Amino acid import rates measured in K562 SLC7A5 and NTC CRISPRi in RPMI and in low leucine medium for SLC7A5 substrates. Data determined from the linear fit ± SE of n ≥ 3. Data for other amino acids are in Figure S2I. (g) Cellular consumption rates of K562 SLC7A5 and NTC CRISPRi in RPMI. (h) Consumption of leucine from low leucine medium by K562 SLC7A5 and NTC CRISPRi cells. (g-h) Data determined from linear fit ± SE of n = 6. (i) Intracellular amino acid levels in K562 SLC7A5 and NTC CRISPRi cultured in RPMI and in low leucine medium (n = 8 ± SEM). Data for other amino acids are in S2J.

To test the effect of medium composition on growth phenotypes more broadly, we assayed all CRISPRi/a cell lines targeting the seven genes in the SLC7 family in a medium (PAA-RPMI) formulated to contain all amino acids as well as five other known SLC7 family substrates (citrulline, ornithine, creatine, creatinine, and carnitine) at levels found in human plasma^20^. The composition of PAA-RPMI contrasts with that of RPMI which contains many amino acids at 0.4× to 10× the levels found in human plasma but only trace amounts of alanine, cysteine and the five other SLC7 substrates (**Table S2**). However, in competition assays, we found that RPMI phenotypes were recapitulated in PAA–RPMI (Pearson correlation coefficient = 0.94), suggesting that differences in the levels of amino acids and other substrates between RPMI and human plasma do not substantially influence the phenotypes of SLC7 family hits. We also confirmed the absence of a phenotype for OE of SLC7A8 (LAT2) despite RT-qPCR confirmation that SLC7A8 levels had been significantly upregulated (**Figure S2C**). SLC7A8 is annotated as an importer for neutral amino acids such as Leu, Ile, His, and Phe^12,33^, but our findings suggest that SLC7A8 expression does not result in import of neutral amino acids under growth-limiting concentrations in K562 cells (this conclusion is also consistent with the absence of a CRISPRi phenotype for SLC7A8 under all screening conditions).

### Knockdown and overexpression by CRISPRi/a directly changes amino acid transport rates

We next tested whether hits obtained in screens reflect changes in the rate of transport of the limiting amino acid caused directly by a change in expression of the gene targeted by CRISPRi/a. We quantified amino acid import by incubating K562 cell lines for different periods of time (seconds to minutes) in medium containing 16 amino acids labeled with heavy isotopes followed by measuring intracellular isotope accumulation using gas chromatography-mass spectrometry (GC-MS). In this approach, separate dishes of cells were used for each time point and import rates for all 16 amino acids were determined from the slope of the increase in labeled amino acid in cell extracts at early time points (**Methods**; **Figure 2B and S2D**). Since many transporters function as exchangers, most amino acids are both imported into and exported out of cells^10,12^. The balance between import and export represents the net transport across the plasma membrane and can be determined by measuring the consumption of amino acids in the culture medium (**Figure S2D**). We quantified import and net transport in K562 cells in complete medium and determined absolute rates using an external standard curve for all 20 amino acids analyzed by GC-MS in the same conditions^21^ (**Figure 2C and S2E-G**).

We found that, while most amino acids were consumed by cells and depleted from the growth medium, five amino acids (Asn, Asp, Glu, Gly, Pro) underwent net secretion, consistent with previous findings in non-small cell lung cancer lines^3^. These data are also consistent with our finding that removal of Asn, Asp, Glu, Gly or Pro from RPMI did not reduce proliferation of K562 cells (**Figure 1C, 2C**). For nine out of the 11 amino acids consumed by K562 cells, import rates exceeded net transport rates by >7-fold, demonstrating high bidirectional flux (**Figure S2H**). We also measured intracellular free amino acid levels and extrapolated free amino acid pool turnover rates by dividing import rates by intracellular levels. We found that most amino acid pools were turned over rapidly (in minutes), further demonstrating the high flux of amino acids in cells (**Figure 2D**).

Using these transport assays, we measured the effects of KD of SLC7A5 on amino acid transport rates; SLC7A5 is a protein required for tumor growth in many settings^35^. In screens, CRISPRi of SLC7A5 did not reduce proliferation of K562 cells in complete RPMI but induced a strong defect in a medium low in any one of the neutral amino acids His, Ile, Leu, Met, Phe, Trp, Tyr, or Val (**Figure 2E**). When we measured amino acid import into SLC7A5 KD cells growing in either RPMI or low Leu conditions, we observed a large reduction in the rates of import of these neutral amino acids (**Figure 2F**), whereas import rates for all other amino acids were unchanged (**Figure S2I**). Consistent with an absence of proliferation phenotype for SLC7A5 KD in RPMI, amino acid consumption rates were unchanged (**Figure 2G**). Despite a significant reduction in neutral amino acid transport caused by SLC7A5 KD, import rates still exceeded consumption rates (**Figure 2C,F,G**). Thus, a strong transport phenotype does not necessarily result in a growth phenotype.

In low Leu medium, however, Leu import rates in control cells were below RPMI consumption rates and import rates were even lower in SLC7A5 KD cells, consistent with a reduction in proliferation imposed by the low amino acid medium and the strong phenotype of SLC7A5 KD in that medium (**Figure 2F,G**). More specifically, measured Leu consumption rates in low Leu medium were 6-fold lower in control cells and 14-fold lower in SLC7A5 KD cells as compared to Leu consumption in RPMI (**Figure 2H**). Moreover, reduced import rates for SLC7A5 substrates were generally associated with reduced free intracellular amino acid levels in both RPMI and low Leu (**Figure 2I** and **S2J**). Of note, import rates of SLC7A5 substrates other than Leu were higher in low Leu conditions than in RPMI (**Figure 2F** and **S2K**), consistent with a model in which multiple substrates are competing for import by a single transporter. Moreover, intracellular levels of amino acids other than Leu were generally higher in low Leu medium as compared to RPMI. This likely arises because proliferation rates were lower and import rates for amino acids other than Leu were either unchanged or higher (**Figure S2L**). Overall, these results show that KD of SLC7A5 reduces the import rate of all of its amino acid substrates, and that this reduction has an effect on cell proliferation specifically when transport rates of one amino acid falls below the consumption rates. These and similar findings strongly suggest that many phenotypes in our CRISPRi/a screens reflect direct changes to the rate of transport of the limiting amino acid by the targeted transporter gene.

### SLC7A6 and SLC7A7 can scavenge cationic amino acids

We next used transport assays to characterize phenotypes associated with SLC7A6 and SLC7A7 perturbations (**Figure 2A, 3A**). It is known that whereas SLC7A1, SLC7A2, and SLC7A3 use the cell membrane potential to drive import of Lys and Arg, SLC7A6 and SLC7A7 exchange Arg or Lys for a neutral amino acid (+ Na^+^)^33^. In physiological conditions with ongoing import of cationic amino acids via SLC7A1–3, it is thought that both SLC7A6 and SLC7A7 concentrate neutral amino acids by net exporting Arg and Lys^12,33^. However, we found that OE of SLC7A7 resulted in a strong proliferation advantage in low Lys medium and OE of SLC7A6 conferred a proliferation advantage in both low Lys and low Arg media, suggesting that these proteins may also act as net importers of Arg and/or Lys in some conditions. Supporting this hypothesis, we found OE of SLC7A7 in cells grown in RPMI increased Lys and Arg import to almost the same degree as OE of SLC7A1 (4–8-fold), which is the main importer of these amino acids in K562 cells (**Figure 3B**). However, whereas SLC7A1 OE increased intracellular Arg and Lys levels, SLC7A7 OE increased the levels of multiple large neutral amino acids, such as Leu, Ile, Phe, Val, and Tyr but did not result in a net increase in intracellular Arg and Lys levels (**Figure S3A**). These results highlight the difference in transport mechanism by SLC7A7 (exchanger) compared to SLC7A1 (importer). SLC7A1 OE results in Arg and Lys accumulation in cells and confers a fitness advantage in low Arg and low Lys conditions. In contrast, SLC7A7 OE increases import of both Lys and Arg but does not lead to intracellular accumulation due to SLC7A7 being an exchanger. The growth phenotype of SLC7A7 OE reflects the balance of transport in specific conditions: net import of Lys in low Lys and absence of net import of Arg in low Arg. By extension, we hypothesize that SLC7A6 OE increases the net import of both Arg and Lys in conditions where either is limiting proliferation.

**Figure 3.**
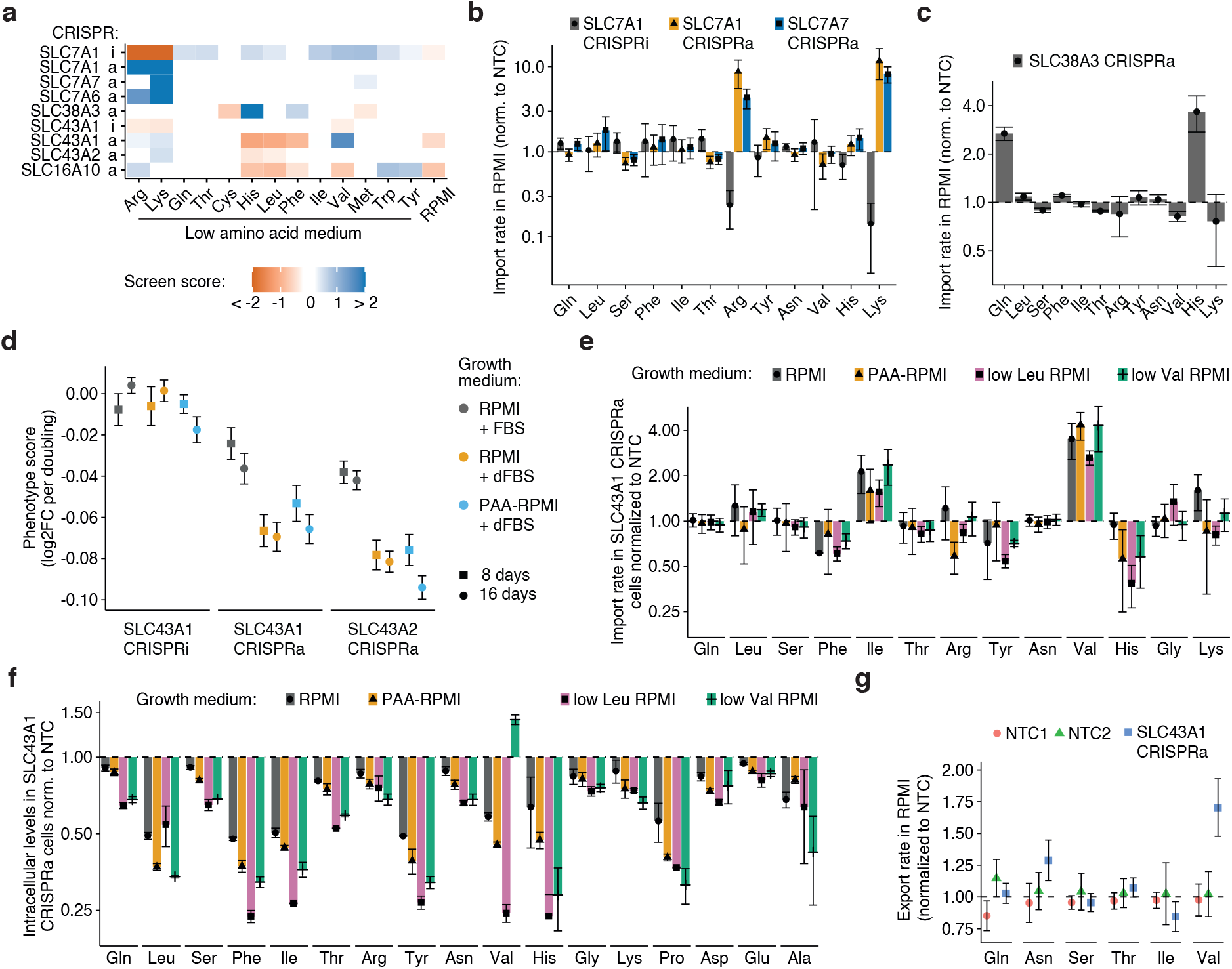
SLC43A1/LAT3 is a net exporter of large neutral amino acids. (a) Scores obtained in transporter CRISPRi/a screens in low amino acid medium and in RPMI. (b) SLC7A1 CRISPRi/a and SLC7A7 CRISPRa specifically alter the import of arginine and lysine in K562 cells in RPMI. (c) SLC38A3 CRISPRa increases import of histidine and glutamine into K562 cells in RPMI. (d) CRISPRa of SLC43A1 and SLC43A2 induces a proliferation defect in K562. Phenotype scores were determined in competition assays and represent the average log2 fold change between a specific and a non-targeting control (NTC) sgRNA normalized by the number of population doublings. Conditions are RPMI with regular fetal bovine serum (FBS), RPMI with dialyzed FBS (dFBS), and RPMI modified such that amino acids match human plasma levels (PAA–RPMI). Data are the mean ± SEM of 2 biological replicates each with 6 technical replicates. (e) CRISPRa of SLC43A1 increases the import rate of isoleucine and valine into K562 cells. (f) CRISPRa of SLC43A1 decreases the intracellular levels of large neutral amino acids in K562 cells. Levels were determined from import assays in (e) (n = 7 ± SEM). (g) CRISPRa of SLC43A1 increases the export rate of valine from K562 cells cultured in RPMI. Rates were determined from the linear fit ± SE of n ≥ 3 and normalized to NTC. (b,c,e) Import rates were determined from the linear fit ± SE of n ≥ 3 and were normalized to NTC.

### SNAT3 is a high affinity histidine transporter

SLC38A3 (SNAT3) can transport His and Asn, but is primarily considered to be a Gln transporter, resulting in net import or export depending on environmental conditions^36^. However, we found that OE of SLC38A3 conferred a proliferation advantage in low His but not in low Gln conditions (**Figure 3A**). When we measured amino acid import in RPMI, we observed an increase specifically for His and Gln (**Figure 3C** and **S3B**). These data suggest that SLC38A3 functions as a high affinity His transporter and a low affinity Gln transporter in K562 cells.

### SLC43A1 (LAT3) is a net exporter of large neutral amino acids

In addition to SLC7A5 and SLC7A8, the L–type amino acid transporter (LAT) family includes SLC43A1 (LAT3) and SLC43A2 (LAT4), proteins that are annotated as low affinity facilitated diffusers of the neutral amino acids Leu, Phe, Ile, Met, and Val^10,37,38^. We found that OE of either SLC43A1 or SLC43A2 reduced proliferation rates in RPMI, which is unexpected for an amino acid transporter (**Figure 3A**). OE phenotypes in low amino acid conditions were also unexpected; SLC43A1 OE conferred resistance to low Val but hypersensitivity to low His, Leu, and Phe, suggesting different affinities for the structurally similar amino acids Val and Leu. To investigate these phenotypes further we generated SLC43A1 and SLC43A2 OE cell lines and performed transport assays (**Figure S3C**).

We first confirmed the proliferation defect of SLC43A1 or SLC43A2 OE using competition assays in RPMI and then in other complete media conditions (**Figure 3D**). We then measured amino acid import in RPMI, PAA–RPMI, low Leu, and low Val media and found that import rates in both control and SLC43A1 OE cells correlated with the level of that amino acid in the medium (**Figure S3D,E**). We also observed that OE of SLC43A1 caused a 1.6–2.4-fold increase in Ile import, a 2.6–4.4-fold increase in Val import, but little change in transport of other amino acids across all four types of media (**Figure 3E**). Despite an increase in import rates, intracellular levels of Ile and Val were significantly reduced under these conditions (**Figure 3F**). In addition, the intracellular levels of other neutral amino acids such as Leu, Phe, Tyr and His were reduced by at least 50% in all four media (**Figure 3F**). The only exception to the observation that OE of SLC43A1 leads to a general decrease in neutral amino acid levels was an increase in intracellular Val in low Val conditions, consistent with the resistance phenotype observed in screening data (**Figure 3A,F**). SLC43A2 OE elicited similar changes to import and intracellular levels in RPMI as SLC43A1 OE (**Figure S3F,G**). To explain these phenotypes, we hypothesized that SLC43A1 and SLC43A2 are net exporters of neutral amino acids such as Leu, Phe, Ile, Tyr, Val, and His, in most conditions and net importers of Val and Ile only in specific conditions.

To test this, we measured amino acid export by incubating cells in RPMI containing heavy-isotope labeled amino acids for an extended period of time (>2 hr), rapidly switching the medium to regular RPMI, and then measuring release of labeled amino acids into the medium. Despite being technically challenging due to the rapidity of export and the confounding effects of differences in intracellular amino acid levels between cell lines on export rates, we were able to quantify export rates for a subset of amino acids. We found that export of Val, but not Gln, Asn, Ser, or Thr, was increased upon SLC43A1 OE (**Figure 3G**). Despite reduced intracellular levels, export of Leu, Ile, and Phe in SLC43A1 OE was similar to control, suggesting a higher intrinsic export capacity (**Figure S3H**). In aggregate these data suggest that SLC43A1 and SLC43A2 are amino acid exchangers that preferentially facilitate neutral amino acid export, but can also import Ile and Val in some conditions. The balance of import to export is condition-specific as measured either by the effect of transporter KD/OE on growth or in transport assays. These results (i) demonstrate the role of the environment in modulating transport, (ii) illustrate the phenotypic complexity that arises from changes in bidirectional transport, and (iii) highlight the importance of directly quantifying both import and export flux in determining net transporter activity.

### Serotonin acts as an endogenous antioxidant to prevent ferroptosis

CRISPRi/a screens in low amino acid conditions have the potential to inform on indirect effects induced by substrates that alter the ability to transport the limiting amino acid, change consumption or demand for a limiting amino acid, or affect cell viability in a low amino acid environment. For example, OE of the Gly transporter SLC6A9 caused hypersensitivity to low Ser, consistent with the finding that conversion of Gly to Ser leads to depletion of the one-carbon pool in these conditions (**Figure 1G**)^39^.

Among the low amino acid conditions we examined in K562 cells, low Cys was unique because it caused high cell death (**Figure 1C**), likely via cysteine and glutathione depletion inducing ferroptosis^30,31^. We hypothesized that transporter screens in low Cys media would yield hits related to transport of cystine (the oxidized conjugate of cysteine most abundant in culture medium), but also of other molecules that might alter sensitivity to ferroptosis. Consistent with this hypothesis we found that one of the strongest hits in low Cys screens was SLC7A11 (**Figure 4A**), the main importer of cystine in animal cells^40^. Since ferroptosis is dependent on iron availability, KD of multiple iron transporters also affected cell viability; KD of SLC11A2 (DMT1), the main importer of iron, or SLC25A37 (Mitoferrin–1), a mitochondrial transporter of iron, conferred resistance to low Cys, and KD of SLC40A1 (Ferroportin), the main exporter of iron^10,41^, lead to hypersensitivity (**Figure 4A**). KD of ABCC1 (MRP1), one of the three major multidrug efflux pumps, also conferred resistance to low Cys, likely by lowering the efflux of intracellular glutathione^42^, suggesting that multidrug resistance via ABCC1 OE might induce sensitivity to ferroptosis, a potentially druggable vulnerability.

**Figure 4.**
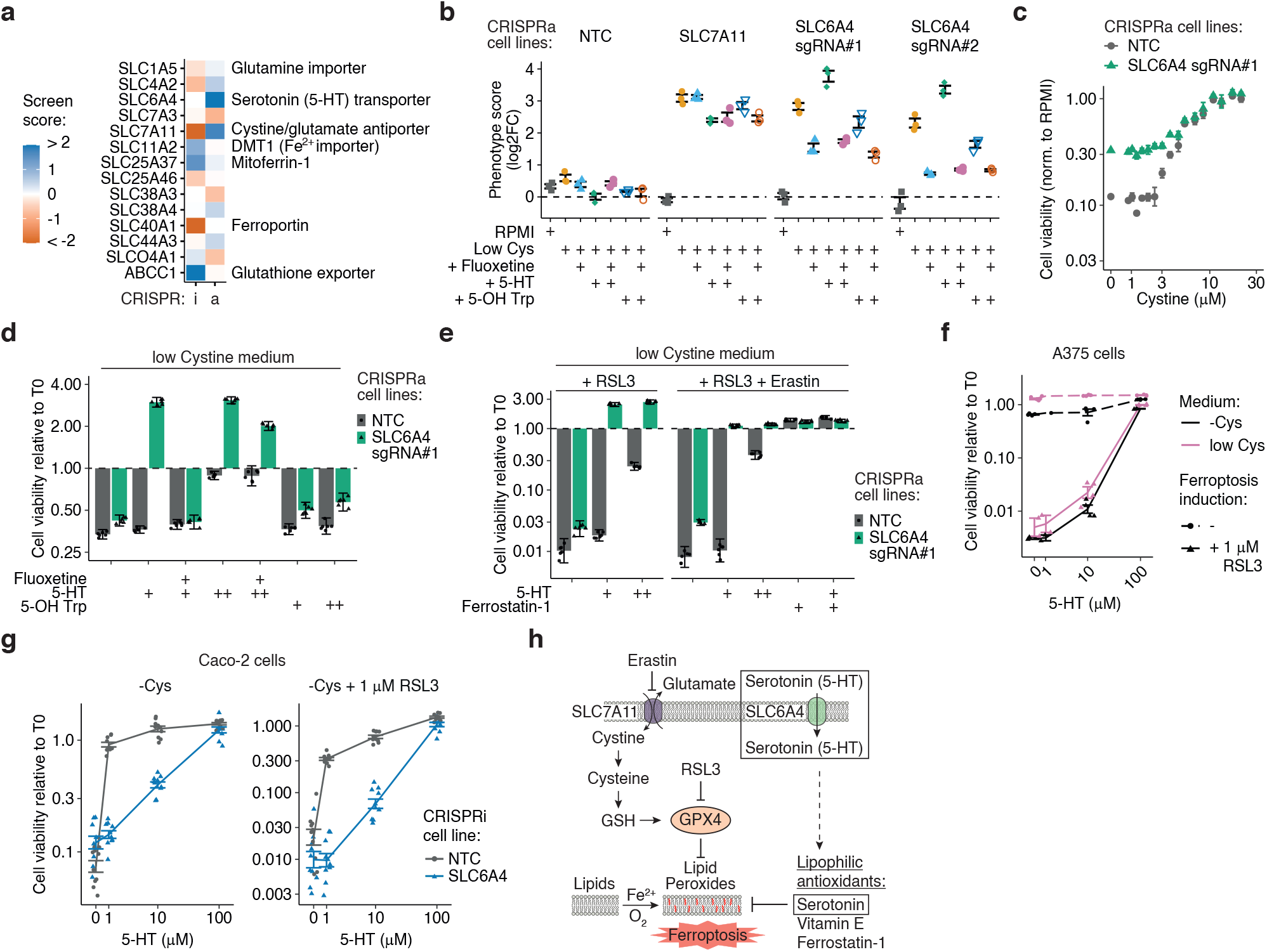
Serotonin (5-HT) protects cells from ferroptosis via its endogenous antioxidant activity. (a) Scores of hits obtained in transporter CRISPRi/a screens in K562 cells in low cystine medium. (b) SLC6A4 CRISPRa provides a growth advantage in low cystine dependent on SLC6A4 activity. Phenotype scores were determined in competition assays in low cystine medium replicating screen conditions and represent the average log2 fold change between a specific and a non-targeting control (NTC) sgRNA. The contribution of SLC6A4 to the phenotype was tested via addition of the SLC6A4 inhibitor, fluoxetine (10 μM), addition of exogenous 5-HT (1 μM), or of 5-hydroxytryptophan (5-OH Trp; 1 μM) the metabolic precursor of 5-HT. Data: n = 3 ± SEM. (c) SLC6A4 CRISPRa provides resistance to K562 cells over a range of low cystine levels. Cell viability was quantified in a luminescent assay after 48 hr in low cystine (n = 4 ± 95% confidence interval). (d) 5-HT protects K562 cells from death in low cystine medium and expression of SLC6A4 greatly increases protection. Assay as in c) and data (n = 6 ± SD) were normalized to the initial cell count (T0). Fluoxetine: 5 μM; 5-HT: 1 (+) or 10 (++) μM; 5-OH Trp: 1 (+) or 10 (++) μM. (e) same as (d). Ferrostatin-1: 0.1 μM; RSL3: 1 μM; Erastin: 5 μM. (f) 5-HT protects A375 cells from ferroptosis only at high concentration. Assay as in d). Data: n = 4 ± SEM. (g) 5-HT protects Caco-2 cells from ferroptosis dependent on SLC6A4 expression and independent of GPX4 activity. Assay as in d). Data: n = 10 ± SEM. (h) Cartoon illustrating the role of 5-HT and SLC6A4 in suppressing ferroptosis. Boxed areas are from this study and core ferroptosis pathway components and ferroptosis-modulating molecules are adapted from others^31,66^.

Unexpectedly, we found that OE of SLC6A4, the serotonin (5-hydroxytryptamine; 5-HT) transporter, also conferred resistance to low Cys (**Figure 4A**). SLC6A4 transports the neurotransmitter 5-HT from the synaptic cleft back into pre-synaptic neurons and is the target of selective serotonin reuptake inhibitors (SSRIs), a class of anti-depressant drugs^43^. SLC6A4 is poorly expressed in K562 cells whereas SLC7A11 is well expressed, and RT-qPCR confirmed that CRISPRa induced strong and specific OE of SLC6A4 or SLC7A11 (**Figure S4A**). Competition assays in screening conditions performed over a 14-day period confirmed that SLC6A4 and SLC7A11 OE cells were resistant to low Cys conditions (**Figure 4B**). When we added the SSRI fluoxetine to low Cys medium to block 5-HT transport by SLC6A4, resistance of SLC6A4 OE cells to low Cys was reduced by at least 50%, and when we added excess 5-HT, resistance was increased (**Figure 4B**). We also tested OE cells in viability assays in low Cys over a 72 hr period in the absence of media changes and again saw that OE of SLC6A4 induced resistance to low Cys (**Figure 4C,D**). This effect could be augmented by addition of exogenous 5-HT and reduced by addition of an SSRI (**Figure 4D**). Thus, transport of 5-HT into cells by SLC6A4 appears to be the cause of ferroptosis resistance. While not part of the RPMI formula used in the low Cys screens, 5-HT is potentially present in serum and as a contaminant in commercial preparations of Trp; it can also be generated via oxidative degradation of Trp in solution^44^. Addition to media of 5-hydroxytryptophan (5-OH Trp), the metabolic intermediate in the 2-step conversion of Trp into 5-HT, had no effect on viability, suggesting that it either did not protect cells from ferroptosis or that it was not imported (**Figure 4B,D**). Based on these data, we hypothesize that SLC6A4 OE confers a growth advantage in low Cys conditions due to increased transport of the low level of 5-HT naturally present in the culture medium and a consequent inhibition of ferroptosis.

To identify the steps in the ferroptosis pathway affected by 5-HT, we tested the effect of two ferroptosis inducers: erastin, which inhibits SLC7A11, and RSL3, which inhibits glutathione peroxidase 4 (GPX4). We found that exposure of cells to RSL3 alone or RSL3 plus erastin in low Cys medium did not prevent rescue of ferroptosis by 5-HT present in the medium or added exogenously, suggesting that 5-HT acts downstream of GPX4 (**Figure 4E**). The indole ring of 5-HT makes it a strong chemical antioxidant with a high affinity towards lipid bilayers, especially those containing unsaturated lipids^45,46^. Thus, 5-HT may prevent ferroptosis by acting directly as a radical-trapping antioxidant.

K562 cells or A375 melanoma cells normally express very low endogenous levels of SLC6A4. This likely explains why ferroptosis resistance requires either OE of SLC6A4 or the addition of high concentrations of 5-HT to the medium (**Figure 4D–F** and **S4A**). We also examined ferroptosis protection by 5-HT/SLC6A4 in the intestinal cell line Caco-2, which expresses SLC6A4 at high levels (**Figure S4A**). In Caco-2 cells, addition of 5-HT to low Cys medium prevented cell death, and KD of SLC6A4 blocked that rescue (**Figure 4G**). However, high concentrations of 5-HT in media provided rescue independent of SLC6A4 expression, presumably due to other import mechanisms. Moreover, protection from ferroptosis by 5-HT occurred downstream of GPX4 as demonstrated by experiments with RSL3. Overall, these data show that 5-HT inhibits ferroptosis and expression of SLC6A4 increases the magnitude of inhibition (**Figure 4H**). Importantly, inhibitors of SLC6A4, such as SSRIs, obviate the protection afforded by increased 5-HT levels and might therefore have a role in promoting ferroptosis in SLC6A4-expressing cancers.

### Transporter essentiality is highly condition-specific and can be used to decipher transporter function

To complement CRISPRi/a screens performed in growth-limiting media we performed screens under more physiological conditions in which growth was not deliberately nutrient-limited. We hypothesized that this might yield condition-specific transporter phenotypes that could be used to study intracellular metabolism and the composition of the environment, as transport flux is influenced by cellular demands and extracellular nutrient levels. First, we performed CRISPRi screens in K562 cells cultured in complete RPMI and found that about 9% of all transporter KD were significantly depleted (representing 46 essential genes), with results highly reproducible between biological replicates (**Figure 5A,B** and **S5A-C**). We also explored transporter essentiality in three other conditions: (i) RPMI at high confluence, (ii) RPMI with regular fetal bovine serum (FBS) replaced with FBS that was dialyzed to remove small molecules (dFBS), and (iii) Dulbecco’s Modified Eagle Medium (DMEM), an alternative synthetic medium.

**Figure 5.**
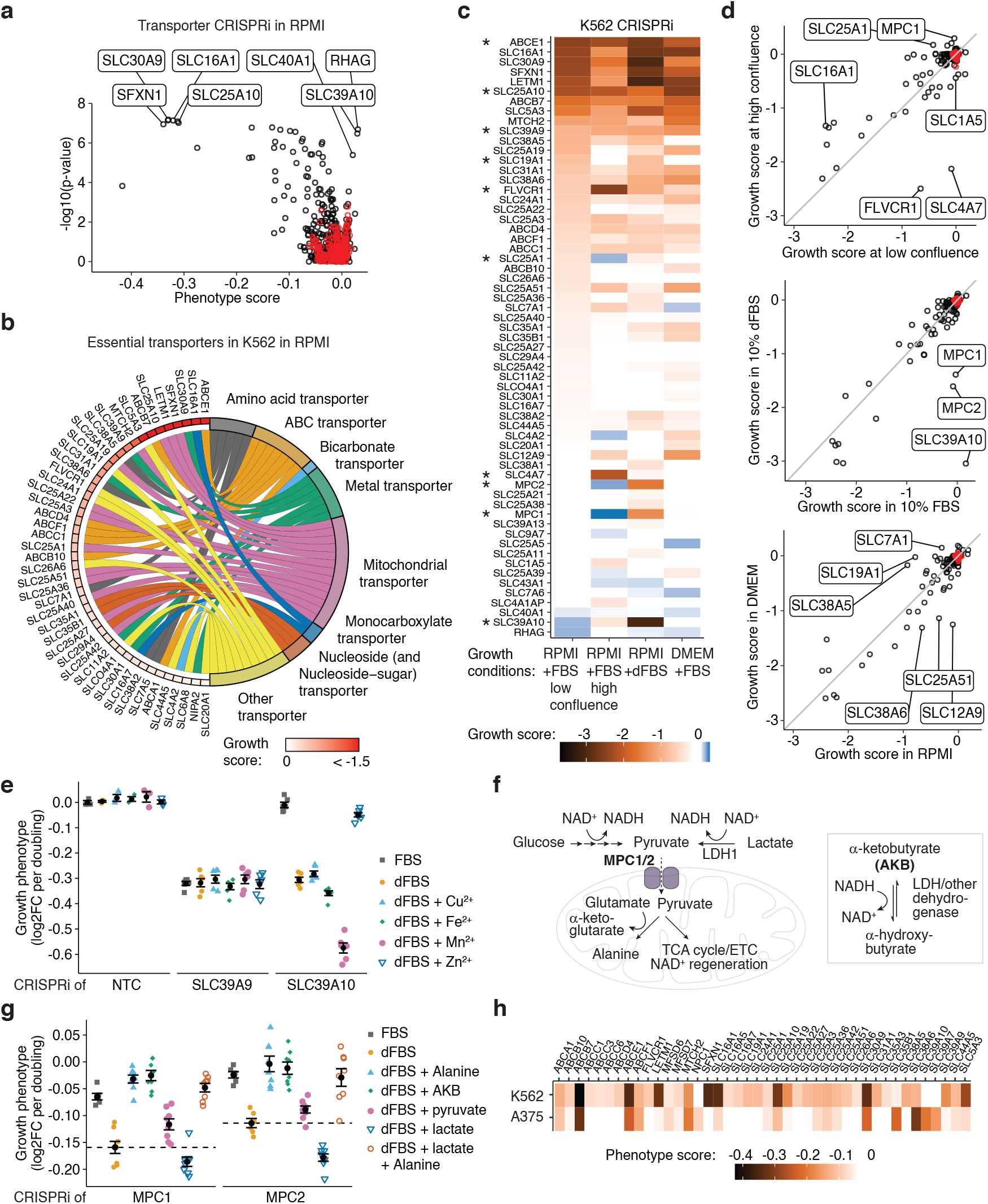
Transporter essentiality is highly condition-specific and can be leveraged to decipher transporter function. (a) Volcano plot of transporter CRISPRi screens in K562 cells growing in RPMI. Black circles represent transporter genes and red circles represent negative control genes. Data were computed from 2 screens each with 2 technical replicates. (b) Chord plot of all essential transporters in K562 cells growing in RPMI. Growth scores were determined from (a). (c) Transporter essentiality in K562 cells is highly dependent on the environment. Tile plot of growth scores determined in CRISPRi screens in K562 cells in various growth conditions (n = 2 replicates). All transporters significantly enriched or depleted in at least one condition are included. Asterisks highlight transporters discussed in the main text. (d) Comparison of essentiality in two conditions identifies transporters most affected by the environmental change. Data represent growth scores for all transporters as determined in (c) (Black circles = transporters; red circles = negative control genes). (e) Growth complementation assays identify SLC39A10 as the main zinc importer in K562 cells and manganese as a competing ion. Phenotypes determined in competition assays of K562 CRISPRi cell lines growing in RPMI + FBS, in RPMI + dialyzed FBS (dFBS), or in RPMI + dFBS supplemented with 10 μM metal ion. Data represent mean ± SEM of 3 technical replicates for NTC and 3 technical replicates of 2 biological replicates for SLC39A9 and SLC39A10. (f) Cartoon illustrating the role of the mitochondrial pyruvate carrier (MPC1/2) in shuttling pyruvate into the mitochondria, and the major sources and uses of cytoplasmic and mitochondrial pyruvate. Exogenous alpha-ketobutyrate (AKB) is used in this study to restore NAD^+^ levels. (g) Addition of either alanine or of molecules restoring NAD^+^ levels alleviate the defect of K562 MPC1/2 CRISPRi growing in RPMI + dFBS. Phenotypes determined in competition assays of K562 MPC1 or MPC2 CRISPRi cell lines in RPMI + FBS and in RPMI + dFBS with 130 μM alanine, 1 mM AKB, 1 mM pyruvate, 4.5 mM lactate, or 130 μM alanine + 4.5 mM lactate. Data: mean ± SEM of 2 biological replicates each with 3 technical replicates. (h) Comparison of transporter essentiality in K562 and A375 cells. Data (n = 2 replicates) are the phenotype scores determined in CRISPRi screens in RPMI for all transporters significantly depleted in at least one cell line.

We found that a minority of KD phenotypes (e.g. ABCE1, SLC25A10, and SLC39A9) were of similar magnitude across all conditions (**Figure 5C,D**), suggesting that extracellular nutrient levels do not have a major effect on the biological roles of these proteins. The main function of ABCE1 is not as transporter but in assisting translation reinitiation and ribosome recycling^47^. SLC25A10 is a mitochondrial dicarboxylate transporter, and SLC39A9 is localized to the Golgi apparatus and plays a role in intracellular zinc trafficking^26,48^. In contrast, the great majority of transporters exhibited strong condition-specific essentiality (**Figure 5C**). For example, in a highly confluent environment, cells experience worse proliferation defects following KD of the choline importer FLVCR1 (also known as SLC49A1) and the bicarbonate transporter SLC4A7, and a growth advantage for KD of the mitochondrial citrate transporter SLC25A1^26,49^. Further, the folate transporter SLC19A1 was only essential in RPMI at low confluence or with dFBS.

To identify the nutrients underlying differential phenotypes, we used growth complementation assays where specific nutrients were added to environments with proliferation defects to attempt to rescue that defect. For example, a comparison of essentiality in K562 cells cultured in RPMI + dFBS to essentiality of cells grown in complete RPMI identified three transporters (SLC39A10, MPC1 and MPC2) with strongly divergent phenotypes (**Figure 5D**). KD of SLC39A10 conferred a strong proliferation defect in RPMI + dFBS but not in RPMI + FBS, whereas KD of the closely related gene SLC39A9 was equally deleterious in both conditions. The SLC39 family contains 14 genes that are annotated as zinc importers, although a subset of family members import metal ions such as cadmium and manganese^48^. Dialysis removes small molecules and metal ions not bound to protein, and RPMI contains no transition metals. We therefore hypothesized that SLC39A10 is the main importer of one or more metal ions essential to K562 cells. In contrast, SLC39A9 is buffered from extracellular changes in metal ion concentration because of the protein’s localization and function in the Golgi^48^. We measured the proliferation of K562 SLC39A9 and SLC39A10 KD cells in assays in which specific metal ions were added back to RPMI + dFBS (**Figure 5E** and **S5D**). Consistent with screening results, KD of SLC39A9 reduced proliferation across all conditions, but KD of SLC39A10 was deleterious only in RPMI + dFBS. Moreover, addition of 10 μM zinc to RPMI + dFBS rescued the proliferation defect of SLC39A10 KD and addition of manganese exacerbated it, suggesting that SLC39A10 is an important zinc importer in K562 cells and manganese can compete for import (**Figure 5E**).

KD of the mitochondrial pyruvate carrier (MPC) was also selectively deleterious in RPMI + dFBS. The MPC is composed of subunits MPC1 and MPC2 and shuttles cytoplasmic pyruvate derived from glucose into the mitochondria (**Figure 5F**). MPC knockouts are deleterious in low alanine conditions because mitochondrial pyruvate is required to synthesize alanine^50^. We measured the growth phenotype of K562 MPC1 and MPC2 KD cell lines in competition assays and found that both cell lines exhibited a strong proliferation defect in RPMI + dFBS but not in RPMI + FBS; moreover, the defect observed in RPMI + dFBS was fully rescued by addition of alanine, an amino acid that is absent from RPMI (**Figure 5G and S5E**). As lactate and pyruvate are also present in FBS and removed by dialysis (**Figure S5F**), we tested the effects of adding these to MPC KD cells cultured in RPMI + dFBS; we found that lactate exacerbated the proliferation defects of MPC1 or MPC2 KD and pyruvate alleviated them (**Figure 5G** and **S5G**). Moreover, media supplementation with either pyruvate or alanine fully rescued proliferation defects caused by lactate addition to MPC1 or MPC2 KD cells (**Figure 5G** and **S5G**). Based on the differing effects of lactate and pyruvate on proliferation and their known influence on the NAD^+^/NADH ratio, we hypothesized that lower proliferation rates were caused by a reduction in the NAD^+^/NADH ratio^51^. To test this we supplemented RPMI + dFBS with α-ketobutyrate (AKB), a molecule that increases the NAD^+^/NADH ratio without contributing carbon atoms to the tricarboxylic acid (TCA) cycle or generating ATP^52^; we observed that the proliferation defects caused by MPC KD were fully alleviated (**Figure 5G**). These results are consistent with a requirement for mitochondrial pyruvate to synthesize alanine and to regenerate NAD^+^ via the electron transport chain. Cell growth in conditions that alleviate one or both of those requirements reduces dependence on MPC activity, and thus mitigates the effect of MPC KD on proliferation. In sum, we were able to trace down the condition-specific essentiality of MPC to differences in alanine and pyruvate levels in various media.

We also started to explore the effects of cell type on the requirement for specific transporters by comparing transporter essentialities in A375 (BRAF^V600E^ melanoma) and K562 cells. A375 cells are advantageous for this purpose because sgRNAs can be introduced efficiently, CRISPRi/a is highly effective, and they can be cultured in the same medium as K562 cells^53^. We mitigated potential differences in sgRNA activity across cell lines by comparing average growth phenotypes of a subset of sgRNAs (**Methods**). We found that many transporters exhibited similar essentiality in the two cell lines but that the phenotypes of 17 transporters were strongly cell-line specific (**Figure 5H**). For example, KD of the mitochondrial serine transporter SFXN1^54^ was highly deleterious in K562 cells but had no effect in A375 cells. We confirmed that SFXN1 was expressed in both cell lines and that KD was similarly effective in the two lines (**Figure S5H**), indicating a cell-specific difference in metabolism. This example suggests that comparisons of transporter essentiality between cell lines could be informative in combination with studies on differences in intracellular metabolism. Cell type-specific requirements for transporters may also create opportunities for achieving therapeutic selectivity with transporter inhibitors.

### Transporter CRISPRi/a screens in subcutaneous tumors reveal effects of the tumor microenvironment on metabolism

Given that transporters are highly sensitive to the growth environment, we sought to perturb transporter expression in the setting of actual — albeit immune deficient — xenografted tumors. We therefore generated subcutaneous K562 and A375 tumors in immunodeficient NOD scid gamma (NSG) mice. CRISPRi and CRISPRa transporter libraries were introduced into K562 or A375 cells, the cells were injected subcutaneously in the flank of NSG mice, and sgRNA enrichment/depletion was evaluated by 14 days later. To ensure sufficient library complexity for these in vivo screens, we first optimized injection conditions by determining engraftment efficiencies using GFP-labeled cells (**Figure S6A,B**). In all screens we identified transporter perturbations that lead to changes in proliferation in the subcutaneous tumor environment (**Figure 6A,B**). To identify effects that were unique to the in vivo setting, we also screened the prepared pools of cells in RPMI, in human plasma-like medium (HPLM)^20^, a medium that mimics the nutrients found in human blood, and in adult bovine serum (ABS), another physiological culture medium with blood nutrient levels^40^ (**Figure 6C,D**).

**Figure 6.**
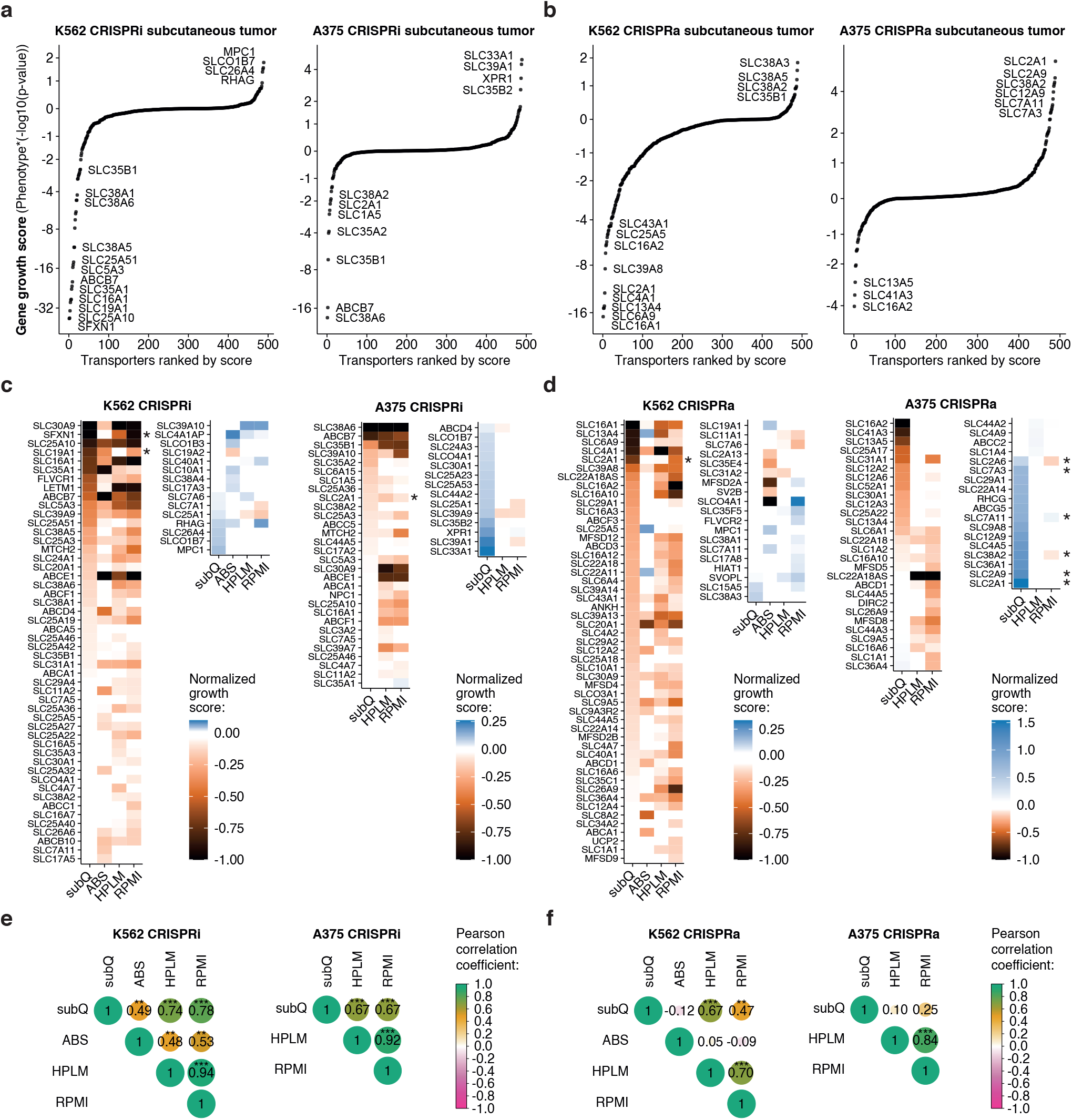
Identification of essential transporters and environmental growth limitations in subcutaneous tumors. (a,b) Pools of K562 and A375 transporter CRISPRi/a libraries were injected subcutaneously in the flank of immunodeficient mice. Growth scores were determined from enrichment/depletion of sgRNAs in whole tumor homogenates (n = 6) relative to the composition of the injected cells. (c,d) Comparison of transporter CRISPRi/a screens in subcutaneous (subQ) tumors to screens in growth culture media identifies condition-specific effects. Pools of cells prepared in (a,b) were cultured in ABS (100% bovine serum), in HPLM (human plasma-like medium), and in RPMI. Data represent all transporters significantly enriched or depleted in at least one condition, and growth scores determined from 2 replicates were normalized to the most depleted transporter in each screen. Asterisks highlight transporters discussed in the main text. (e,f) Correlation analysis of growth scores obtained in different environments. Pearson correlation coefficients were determined in pairwise comparison of growth scores for all significant enrichments/depletions in (c,d). *: p < 1E-2; **: p < 1E-4; ***: p < 1E-6.

We quantified the similarity in phenotypes across environments by calculating pairwise correlations using all genes that had a deleterious or beneficial phenotype in at least one of the environments tested (**Figure 6E,F**). In CRISPRi screens, we found that HPLM and RPMI profiles were strongly correlated (Pearson’s r > 0.9) and these two conditions were more weakly but similarly correlated to screens performed in xenograft tumors (r = 0.7–0.8). Correlations for CRISPRa were generally lower than for CRISPRi, suggesting a greater dependence on the environment for transporter OE phenotypes. In K562 CRISPRa screens, xenograft scores were more strongly correlated with HPLM (r = 0.7) than with RPMI scores (r = 0.5), which implies that the composition of HPLM more closely mimics the nutrient environment of tumors than RPMI^21^. A375 CRISPRa screens were an outlier in this analysis in that scores for xenograft tumors were not correlated with either HPLM or RPMI (r < 0.25), and we found that scores in ABS were poorly correlated to other conditions.

Overall, these CRISPRi/a screens (**Figure 6C,D**) highlight specific transporters that reveal the influence of the environment on metabolism, and differences in metabolism between cell lines. For example, KD of the main glucose importer, SLC2A1 (GLUT1), was deleterious in A375 cells across all conditions tested and OE increased growth in tumors. In contrast, KD of SLC2A1 in K562 cells had no phenotype, and OE was deleterious in tumors and in RPMI. Despite these differences in phenotypes, we observed similar levels of SLC2A1 expression at baseline in both cell lines and OE was equally effective in upregulating expression (~10-fold) (**Figure S6C**). Thus, differences in SLC2A1 phenotypes between the cell types are likely due to differences in glucose metabolism rather than in SLC2A1 expression or activity per se. In particular, A375 cells appear to have a higher glucose demand compared to K562 cells, consistent with reports that oncogenic BRAF^V600E^ in A375 cells increases the rate of glycolysis^55^.

Our studies with low amino acid media suggest that when OE of a nutrient transporter confers a growth advantage, the level of that nutrient in the environment is often limiting proliferation. For example, the growth advantage conferred by OE of SLC2A1 in A375 xenograft tumors, and also of SLC2A6 and SLC2A9, two other glucose transporters, highlights that glucose availability is likely limiting for proliferation of A375 in a subcutaneous environment (**Figure 6D**). Furthermore, the growth advantage conferred by OE of amino acid transporters SLC7A3, SLC38A2, and SLC7A11 suggests that Arg, Gln, and Cys levels could also be limiting for proliferation of A375 tumors. On the other hand, the absence of any growth advantage conferred by transporter OE in K562 cells suggests that K562 cells in a solid tumor environment do not experience nutrient limitations that can be overcome by transporter OE. Consistent with this finding, KD of the major amino acid transporters in K562 cells identified in this work (SLC1A5, SLC7A1, SLC7A5, SLC7A11) did not induce any significant proliferation defect in tumors. However, there was clear evidence of other nutrient limitations in K562 tumors. For example, OE of SLC6A9, the main glycine importer, induced a growth defect specifically in K562 tumors, reminiscent of what we observed in a low serine environment (**Figure 1G**). We also observed that KD of the folate transporter SLC19A1, as well as the mitochondrial serine transporter SFXN1, were particularly deleterious in K562 tumors. These perturbations are expected to reduce the activity of the one-carbon metabolic pathway or lead to depletion of one-carbon units^54^ and their phenotypes suggest that maintaining the one-carbon pool is particularly critical in the tumor environment. Differences between A375 and K562 in nutrient limitations could reflect differences in nutrient demand or in nutrient concentrations within the two tumor environments; it should be possible to distinguish between these possibilities in future studies.

## DISCUSSION

Efforts are underway to identify nutrients that are essential for tumor growth and leverage this for improved anti-cancer therapy^56^. In this work, we developed a strategy based on CRISPRi/a screening to systematically identify the transporters of nutrients in cell and used this approach to characterize amino acid transport in the leukemia cell line K562. Although the major families of amino acid transporters have been well described as a result of decades of research^10,12^, our work enables the first simultaneous interrogation of the contribution of each of the 64 annotated amino acid transporters to import and export of specific amino acids in cells across different conditions. To understand which transporters are required for or capable of import in nutrient-poor conditions, we performed screening at amino acid concentrations that limit proliferation. As these concentrations of amino acids are generally lower than the levels in plasma or tissue interstitial fluid (TIF)^20,21^, our growth-based screens therefore identified high affinity transporters, and not necessarily all of the low affinity transporters that might also contribute to physiological amino acid homeostasis. However, we were able to explore the role of lower affinity transport mechanisms by testing individual CRISPRi/a cell lines in transport assays, as well as by repeating our screens in nutrient rich and physiological conditions. Overall, we were able to identify one or more SLCs required for the transport of all 13 amino acids whose environmental concentration can become limiting for the growth of K562 cells, and a complementary set of transporters able to import specific amino acids when overexpressed.

A clear finding from our studies is that amino acid transport is characterized by high bidirectional import and export flux at the membrane, consistent with proposed models of cellular amino acid homeostasis^12,13^. Although net transport, the balance of those two fluxes, is generally skewed towards amino acid import to sustain cell proliferation, we found that some transporters such as SLC43A1 and SLC43A2 are net exporters and only act as net importers when their substrates, valine or isoleucine, are limiting for proliferation. This finding is consequential since inhibitors of SLC43A1 are being pursued as anti-cancer drugs under the premise that it acts as an importer^38^. Other transporters, such as SLC16A10 (TAT1), showed similar phenotypes (**Figure 3A**), suggesting that they might also primarily function as net exporters. These findings highlight the importance of assessing net transport based on measurement of both import and export rates as well as the role of the microenvironment in setting the balance between import and export fluxes.

### Consequences of the antioxidant activity of serotonin in ferroptosis

We found that serotonin (5-HT) and its transporter SLC6A4 act as an endogenous antioxidant to protect cells from ferroptosis. Lipid peroxides are known to mediate ferroptosis, but the membrane compartment where lipid peroxidation happens is unclear^31,57^. We show that expression of the plasma membrane transporter SLC6A4 greatly increases the protective effect of 5-HT, showing that transport across the plasma membrane is required for 5-HT to reach its site of action as anti-ferroptotic factor. Levels of 5-HT and expression of SLC6A4 are uneven within the human body, with the highest levels of both generally found in the gut and in the central nervous system (CNS), suggesting that resistance to ferroptosis via the antioxidant effect of serotonin might play a role in those environments^58^. Other environments with elevated 5-HT levels include carcinoid tumors, which metabolize most of their tryptophan into 5-HT^59^. Our results also imply that commonly prescribed SSRIs might increase the ferroptotic sensitivity of cells expressing SLC6A4 and exposed to 5-HT, an insight of potential relevance to the CNS^60^.

### Using transporter perturbations to study transporter function and probe the composition of the microenvironment

About 20–30% of SLCs lack a known substrate^9,18^, and finding conditions where KD of one of these poorly characterized transporters causes a specific proliferation defect can unmask its function. In yeast, only ~35% of genes in the *S. cerevisiae* genome are essential in rich medium, but ~97% are essential in at least one condition when deletions are tested across a very wide range of environments^61^. By analogy, we hypothesize that most expressed transporters will be essential in human cells when tested over a sufficiently wide range of conditions and find evidence for this in the limited set of conditions explored in this study. For example, the poorly characterized amino acid transporters SLC38A5 and SLC38A6 exhibit diverging phenotypes in the commonly used media RPMI and DMEM (**Figure 5D**). As shown here, CRISPRi/a-based transporter screens are amenable to a rapid systematic exploration of synthetic media compositions since they achieve high signal-to-noise and can be performed in individual regular-sized tissue culture flasks.

The TME is among the conditions of greatest interest for CRISPRi/a screening to identify essential transporters. To approximate this, we performed screens in two cell lines grown as subcutaneous xenografts and compared the results to those obtained in commonly used culture media to identify perturbations specific to the TME. We found that transporter phenotypes observed in subcutaneous tumors were highly correlated to phenotypes observed in RPMI or in HPLM, a medium that replicates metabolite levels found in human plasma. Nevertheless, multiple transporters displayed large differences in essentiality between the in vivo and in vitro environments and further studies will be required to understand these effects.

The TME is generally considered nutrient poor^2^ and metabolite levels in TIF differ substantially from plasma levels^21^. However, it is unclear which nutrients, if any, are constraining tumor growth. As shown in this work and reported by others^62,63^, the identification of growth advantages conferred by transporter overexpression pinpoints nutrients that are functionally limiting tumor growth. Using this approach, we found that A375 melanoma cells experience glucose and amino acid limitation in xenograft tumors. K562 cells, in contrast, did not experience any significant limitation in the same environment, highlighting potential differences in nutrient accessibility to individual cells within these tumors or changes in environmental metabolite levels induced by the tumor itself. These results are consistent with the observation that TIF levels in lung cancer and pancreatic ductal adenocarcinoma xenografts are markedly different^21^.

In conclusion, we show that CRISPRi/a screening allows the systematic manipulation of transporter activity to study nutrient uptake and secretion under a wide range of growth conditions in vivo and in vitro. This provides new insights into metabolism and growth constraints imposed by the TME. In addition to nutrients, we note that transporter CRISPRi/a screens are suited to studying both drug transport and the influence of metabolism on drug sensitivity, as most small molecule drugs are actively imported into cells via SLCs and can be exported via ABC transporters^64,65^. We anticipate that others will be able to leverage our approach to explore other aspects of transporter biology.

## ACKNOWLEDGEMENTS

We thank Dr. Alexander Muir, Dr. Nicholas Matheson, Dr. Zhaoqi Li, and members of the Sorger and Vander Heiden laboratories for helpful discussions and technical advice. We acknowledge the HMS Systems Biology Flow Cytometry Core Facility, the Bauer Core Facility, the ICCB-Longwood Screening Facility, and the Dana-Farber Flow Cytometry Core Facility. We thank Jonathan Weissman for plasmids Addgene #60955 and #85969, Didier Trono for plasmid Addgene #12260, Bob Weinberg for plasmid Addgene #8454, David Sabatini for plasmid Addgene #19319, and Charles Gersbach for plasmid Addgene #71236.

## AUTHOR CONTRIBUTIONS

C.C. designed the study and performed experiments. C.C. analyzed data with help from A.M.D.. C.C. and A.M.D. developed and performed cellular transport assays. A.M.D. performed Western blots. B.L.G. assisted with experiments. E.C.L., A.M.B., and C.C. performed mouse screens. M.G.V.H. and P.K.S. supervised the project and obtained funding. C.C., A.M.D., and P.K.S. wrote the paper with input from all co-authors.

## METHODS

### Cell culture and chemicals

Cell lines were from ATCC: K562 (CCL–243), A375 (CRL–1619), C2BBe1 clone of Caco-2 (CRL–2102), HEK293T (CRL–3216). K562 cells were grown in RPMI-1640 (Corning 10–040) supplemented with 10% (v/v) heat inactivated fetal bovine serum (FBS) (Gibco 10438026). HEK293T and A375 cells were grown in Dulbecco’s modified Eagle medium (DMEM) (Corning 10–013) supplemented with 10% (v/v) FBS. Caco-2 cells were grown in EMEM (ATCC 30–2003) supplemented with 10% (v/v) FBS. All cell lines were grown at 37 °C and 5% CO2, and penicillin and streptomycin were added to all growth media to final concentrations of 100 U/mL and 100 μg/mL, respectively (Corning 30–002–CI). Cells were tested for mycoplasma contamination using the MycoAlert mycoplasma detection kit (Lonza LT07-318).

(1S,3R)-RSL3 (Cayman Chemical 19288), Erastin (MedChemExpress HY–15763), Ferrostatin-1 (Selleckchem S7243) were dissolved in DMSO, quality controlled and stored by the ICCB-Longwood screening facility. Fluoxetine HCl (Millipore Sigma F132), Serotonin HCl (Tocris 354750), 5-hydroxytryptophan (Millipore Sigma H9772) were prepared in DMSO. For metal ion complementation assays, ZnSO_4_·7H_2_O (Fluka 96500), CuSO_4_ (VWR VW 3312–2), Fe(NO_3_)_3_·9H_2_O (Sigma F–1143), and MnCl_2_·4H_2_O (Sigma M8530) were dissolved in H_2_O to 100 mM. Lactic acid (Fluka 69775) was adjusted to pH 7 with NaOH and diluted with H_2_O to 0.5 M, and sodium pyruvate (Sigma P2256) and alpha-ketobutyrate (AKB; Sigma K401) were dissolved in H_2_O to 300 mM and 1M, respectively.

### Transporter CRISPRi/a sgRNA library cloning

Transporter libraries include sgRNAs targeting 413 SLC genes as defined by the human genome organization (HUGO) gene nomenclature, 28 atypical SLCs, and 48 ABC transporters (45 as defined by HUGO and 3 atypical). sgRNA sequences were from Horlbeck et al.^32^ and included 10 sgRNAs per gene for all genes, except 37 of them that had 2 transcription start sites and were therefore represented by 20 sgRNAs, and 730 non-targeting control sgRNAs (NTC) (see Supplementary Table S1). The design and cloning of sgRNAs was performed as previously reported^32^ with minor modifications. A single pool of oligonucleotides (12,000) for both CRISPRi and CRISPRa libraries was synthesized by Twist Biosciences. PCR reactions were set up using primers specific to either CRISPRi or CRISPRa sequences using Phusion polymerase (NEB) according to the manufacturer’s instructions and using 3 different conditions to minimize amplification bias (HF buffer, GC buffer, GC buffer + 3% DMSO). Each 50 μL reaction included 4.5 ng template, 0.4 μM primers, and was amplified using 8 cycles of 30 s at 98 °C, 30 s at 60 °C (57 °C for the DMSO-containing reaction), and 10 s at 70 °C preceded with 1 min at 98 °C and followed by 1 min at 72 °C. Two replicate sets of reactions were run, and all CRISPRi and CRISPRa amplified libraries were pooled separately. The amplified library (88 bp) was purified by agarose gel electrophoresis and extracted using the QIAquick gel extraction kit (Qiagen). Purified PCR products were digested with BlpI and BstXI and the digested fragment was purified on a 20% polyacrylamide Tris–borate–EDTA (TBE) gel (Thermo Fisher Scientific). Digested fragments were purified by isopropanol precipitation, and DNA concentrations were quantified by fluorescence using the Qubit dsDNA high sensitivity assay kit (Thermo Fisher Scientific). Plasmid pU6-sgRNA EF1Alpha-puro-T2A-BFP (Addgene #60955) was digested using BlpI and BstXI, purified by agarose gel electrophoresis, and extracted using the QIAquick gel extraction kit (Qiagen). Digested PCR fragments were ligated into the restricted vector at a 1:1 molar ratio using T4 DNA ligase and were purified using the MiniElute PCR purification kit (Qiagen). Purified ligated plasmids were electroporated into MegaX DH10B (Thermo Fisher Scientific) and plated on LB/Amp plates. Library coverage was 10,000× for both libraries and was determined by serial dilution and colony counting. After 15 hr at 30 °C, the lawn of bacteria was scraped off the plates and plasmid DNA was prepared using the Plasmid Plus Maxi kit (Qiagen).

### Lentivirus preparation

HEK293T cells were transfected with the lentiviral plasmid, psPAX2 (Addgene #12260) and pCMV-VSV-G (Addgene #8454) in a 2:2:1 molar ratio using lipofectamine 3000 (Invitrogen) according to the manufacturer’s instructions. The growth medium was replaced 6 hr post-transfection and was then harvested at 24–30 hr and 48–54 hr post-transfection. The two harvested growth medium fractions were pooled, centrifuged at 1,000 *×g* for 10 min, and filtered through a 0.45 μm low-protein binding membrane. Lentivirus containing supernatants were stored at −80 °C.

### Media preparation

To prepare complete RPMI lacking all amino acids, 8.59 g of RPMI w/o amino acids, sodium phosphate (US Biological R8999–04A), 2.00 g sodium bicarbonate (Sigma S6014), and 0.80 g sodium phosphate dibasic (Sigma S0876) were diluted in 945 mL deionized H_2_O. After addition of 100 mL dialyzed fetal bovine serum (dFBS, Gibco, Thermo Fisher Scientific 26400044) and 10 mL of penicillin-streptomycin 100× solution (Corning 30–002–CI) and homogenization, the medium was sterilized via 0.22 μm membrane filtration (RPMI+dFBS w/o amino acids). Amino acids were added to this base medium as needed from stock solutions to RPMI levels or to low amino acid screen concentrations, as described in Supplementary Table S2 and S3. To make complete RPMI used as the control arm in low amino acid screens and experiments, all 19 amino acids were added to complete RPMI w/o amino acids to RPMI levels (RPMI rich).

Complete RPMI with amino acids present at physiological levels (PAA–RPMI) was prepared from RPMI+dFBS w/o amino acids as above, except that 965 mL deionized H_2_O was used. To that, amino acids were added to their concentration in human plasma^20^ using the same stock solutions as for RPMI above. In addition to the amino acids present in RPMI, alanine, cysteine, carnitine, citrulline, creatine, creatinine, N-acetylglycine, and ornithine were also added to their level in human plasma. Quantities added are described in Supplementary Table S2. For low amino acid conditions, the corresponding amino acid was added such that the final concentration in PAA–RPMI matched the concentration used in the CRISPRi/a screens.

To prepare RPMI medium for amino acid import assays (RPMI with 16 amino acids present as heavy isotopes), we first added unlabeled cystine, OH-Pro, Met, and Trp to RPMI+dFBS w/o amino acids. This base medium consisting of complete RPMI – 16 amino acids was then used to prepare heavy-labeled RPMI and RPMI with low Leu or low Val. Stock solutions of heavy-labeled amino acids (Cambridge Isotope Laboratories) were added to RPMI −16 amino acids to their concentration in RPMI or to the screen concentration (for Leu and Val), as outlined in Supplementary Table S2.

To prepare PAA–RPMI for amino acid import assays, we prepared a complete PAA-RPMI base as above but lacking 16 amino acids. We then added heavy-labeled amino acids from the same stocks as RPMI to the concentration in PAA–RPMI as outlined in Supplementary Table S2.

RPMI + FBS and DMEM + FBS were prepared as outlined in “Cell culture” section. Human plasma-like medium (Thermo A4899101) was adjusted with 10% dFBS. Adult bovine serum (ABS; Sigma-Aldrich B9433) thawed at 4 °C was adjusted to 50 μM with Cystine·2HCl (Sigma C6727). After 1 hr at 37 °C to ensure dissolution, ABS was filtered through a 0.2 μm low-protein binding membrane.

### Amino acid titrations

Single amino acid dropout RPMI medium was prepared for each of the 19 amino acids present in RPMI by adding 18 amino acids to complete RPMI w/o amino acids. K562 cells grown in complete RPMI were washed 3× using cycles of centrifugation for 5 min at 300 *×g* and resuspension of the cell pellet in phosphate-buffered saline (PBS, Corning 21–040 CV). The final cell pellet was resuspended in PBS at a final density of 10 mio/mL and was added to single amino acid dropout RPMI or complete RPMI to a final density of 0.1 mio/mL. 30 μL of these cell suspensions were pipetted into the wells of 384-well microplates (Thermo Fisher Scientific 164610). Amino acid stock solutions in H_2_O (as described in Supplementary Table S2) were adjusted to 0.005% Triton X-100 using a 100× stock solution in H_2_O and were added to wells of the microplates using a D300 digital dispenser (Hewlett-Packard). We confirmed that the highest concentration of Triton X-100 (0.33 parts per million) did not affect proliferation of K562 cells. Each amino acid was tested along a two-fold dilution series from 1× to 1/1,024× its concentration in RPMI and including a –amino acid control. Wells on plate edges were filled but not used for any measurements. The drug dispensing arrangement for each amino acid was spatially randomized (and re-organized during data analysis) to minimize bias. Each data point was present in quadruplicates and 3 separate identical plates were prepared for viability measurement after 24, 48, and 72 hr incubation. Assay plates were incubated at 37 °C/5% CO_2_, inside containers humidified by sterile wet gauze. After 24, 48, or 72 hr, plates were removed from incubation and cooled at room temperature for 10 min, before dispensing 30 μL of CellTiter–Glo (CTG) (Promega) (1:1 dilution in PBS) into each well. Following a 10 min incubation at room temperature, luminescence was measured in a plate reader (BioTek Synergy H1). A control plate with K562 cells dispensed in complete RPMI was prepared and luminescence was monitored at the onset of the experiment (T = 0) and after 24, 48, 72 hr incubation in the same conditions to determine K562 doubling rates in RPMI. Each data point was averaged across 4 replicates, divided by luminescence at T=0, was internally normalized within each plate, and was displayed on a log2 scale to represent the total number of population doublings.

### Preparation of CRISPRi/a parental cell lines

To generate a K562 cell line stably expressing dCas9-KRAB (K562 CRISPRi), K562 cells were transduced with lentiviral particles produced using vector pMH0001 (Addgene #85969; which expresses dCas9-BFP-KRAB from a spleen focus forming virus (SFFV) promoter with an upstream ubiquitous chromatin opening element) in the presence of 8 mg/mL polybrene (Sigma). A pure polyclonal population of dCas9-KRAB expressing cells was generated by two rounds of fluorescence activated cell sorting (FACS) gated on the top half of BFP positive cells (BD FACS Aria II). In a third round of FACS sorting, single cells from the top half of BFP positive cells were sorted into microplates to establish monoclonal cell lines. The performance of K562 CRISPRi monoclonal lines in knocking down endogenous genes was evaluated by individually targeting 3 control genes (ST3GAL4, SEL1L, DPH1) and measuring gene expression changes by RT-qPCR, and the best performing monoclonal cell line was selected for all work described in this study. The preparation of monoclonal K562 cells expressing CRISPRa machinery (K562 CRISPRa) and of A375 expressing CRISPRi (polyclonal) and CRISPRa (monoclonal) has been described elsewhere^23,53^.

### Preparation of CRISPRi/a cell lines using sgRNAs targeting individual genes and expression analysis by RT-qPCR

Pairs of complementary synthetic oligonucleotides (Integrated DNA Technologies) forming sgRNA protospacers flanked by BstXI and BlpI restriction sites were annealed and ligated into BstXI/BlpI double digested plasmid pU6-sgRNA EF1Alpha-puro-T2A-BFP (Addgene #60955). Oligonucleotides used to build sgRNA targeting individual genes are listed in Supplementary Table S4. The sequence of all sgRNA expression vectors was confirmed by Sanger sequencing, and lentiviral particles were produced using these vectors as described above (see ‘lentivirus production’). Parental CRISPRi/a cells were infected with individual sgRNA expression vectors by addition of lentivirus supernatant to the culture medium in the presence of 8 μg/mL polybrene. Transduced cells were selected using puromycin (2 μg/mL for K562 cells) starting 48 hr post-transduction and over the course of 7 days with daily addition of the antibiotic. After 24 hr growth in puromycin-free medium, 0.1 mio cells were harvested and total RNA was extracted using the RNeasy Plus Mini kit (Qiagen). cDNA was synthesized from 0.1–0.5 μg total RNA using Superscript IV reverse transcriptase (Invitrogen) and oligo(dT)20 primers (Invitrogen) following the manufacturer’s instructions. Reactions were diluted 2–5 fold with H_2_O and qPCR was performed using PowerUp SYBR Green PCR Master mix (Thermo Fisher Scientific), 2 μL diluted cDNA preparation, and 0.4 μM of primers. All qPCR primers are listed in Supplementary Table S4. To calculate changes in expression level of target genes, all gene specific Ct values were first normalized to the Ct value of a reference gene (GAPDH) to find a ΔCt value. Log2 fold changes (log2FC) in expression were then determined by the difference between the ΔCt value of targeting sgRNAs and that of a non-targeting negative control sgRNA (ΔΔCt).

For Caco-2 cells, we used a one-vector CRISPRi system (Addgene #71236) to create polyclonal knockdown cell lines. Pairs of complementary synthetic oligonucleotides (Integrated DNA Technologies) forming sgRNA protospacers flanked by BsmBI restriction sites were annealed and ligated into BsmBI digested plasmid pLV hU6-sgRNA hUbC-dCas9-KRAB-T2a-Puro. Caco-2 cells were infected using lentiviral supernatant produced from the plasmids above. Transduced cells were selected using EMEM containing puromycin at 2–5 μg/mL over 12 days. Changes in expression level due to CRISPRi were quantified as above.

### Transporter CRISPRi/a screens in low amino acid medium

Lentiviral supernatant was prepared for both the transporter CRISPRi and CRISPRa sgRNA libraries as described above (‘lentivirus preparation’) and was stored at −80 °C. The multiplicity of infection (MOI) of both preparations was determined by titration onto target cell lines and quantification of the percentage of BFP^+^ cells 2–3 days post-transduction by flow cytometry (BD Biosciences LSR II).

For transporter CRISPRi/a screens in low amino acid, K562 CRISPRi (or CRISPRa) parental cells (85–100 mio) were transduced with lentiviral supernatant at a MOI of 0.25–0.3 in 250 mL culture medium + 8 μg/mL polybrene in a 225 cm^2^ cell culture flask (Costar). 24 hr posttransduction, cells were harvested and resuspended in 200 mL fresh medium in 2 × 225 cm^2^ flasks. Starting 48 hr post-transduction, the culture medium was exchanged daily and cells were maintained at 0.5–1.0 × mio/mL in puromycin (1.75 μg/mL) in 400–500 mL. After 6 days in puromycin, the proportion of BFP^+^ cells determined by flow cytometry increased to 94–96% of the fraction of viable cells. After recovery for 1 day in puromycin-free medium, 2 × 10 mio library cells were harvested and stored at −80 °C (T = 0 samples). The remaining cells were grown for 24 hr in RPMI/rich (complete RPMI with all amino acids; see Media preparation). 120 mio cells were harvested by centrifugation at 300 ×*g* for 5 min, were washed twice with 25 mL PBS, and the final cell pellet was resuspended in 4.8 mL PBS. Complete RPMI media with 18 amino acids present at RPMI level and 1 amino acid present at a concentration that limits growth of K562 (see Media preparation) was prepared and 50 mL were added to 150 cm^2^ tissue culture flasks (Falcon 355001) which was preheated and equilibrated to 37 °C/5% CO_2_. Library cells were added to the flasks to a final density of 0.1 mio/mL. For all amino acids except Trp and Cys, screens were conducted over the course of 16 days with 3 cycles of 1 day in RPMI/rich and 4 days in RPMI/low amino acid, followed by a final day in RPMI/rich. The growth medium was changed daily on all flasks using centrifugation of a fraction of the culture and resuspension of the pellet into pre-equilibrated growth medium. For medium change after growth in RPMI/rich, cell pellets were washed twice with PBS before resuspension in low amino acid medium. The amount of culture centrifuged was adjusted such that the final cell density after medium exchange was 0.1 mio/mL. For Trp and Cys, screens were conducted over the course of 16 days and media was changed every 36 hr (1× 24 hr RPMI/rich; 4× pulses of 36 hr RPMI/low amino acid; 1× 36 hr RPMI/rich; 4× pulses of 36 hr RPMI/low amino acid; 1× 36 hr RPMI/rich). The RPMI/rich control arm and the low His, low Arg, and low Lys were run in technical replicates. At the end of the screen, 12–15 mio cells were collected by centrifugation, washed once with PBS and stored at −80 °C. The number of population doublings for each screen is indicated in Supplementary Table S3. Genomic DNA (gDNA) was extracted from cell pellets using the QIAamp DNA Blood Mini Kit according to the manufacturer’s instruction, except that the elution was performed using 10 mM Tris·HCl pH 8.5. Typical yields from 15 mio cells ranged from 90 to 240 μg gDNA. sgRNA barcodes were amplified by PCR using the gDNA from at least 6 mio cells as template and Phusion (NEB M0530) as polymerase. An equimolar mix of primers with stagger regions of different length (CC_LSP_025 to CC_LSP_032_c) was used as forward primer and barcoded index primers (CC_Cri_a_rev1 to CC_Cri_a_rev10) were used as reverse primers. Reactions were composed of 1 × HF buffer, 0.2 mM dNTPs, 0.4 μM forward primer mix, 0.4 μM indexed reverse primer, 0.5 μL Phusion, 1.5 mM MgCl_2_, and 5 μg gDNA in a volume of 50 μL. After 30 s at 98 °C, the reactions were subjected to 23 cycles of 98 °C for 30 s, 62 °C for 30 s and 72 °C for 30 s, and were followed by 72 °C for 5 min. All reactions from each screen were pooled and the amplified PCR product (~240–250 bp) was purified by agarose gel electrophoresis using the QIAquick gel extraction kit (Qiagen). Purified PCR products were quantified by fluorescence using the Qubit dsDNA high sensitivity assay kit (Thermo Fisher Scientific). Individual indexed libraries were mixed in equimolar ratio and were further purified using a QIAquick PCR purification kit (Qiagen). After quantification by qPCR using the NEBnext library quant kit for Illumina (NEB), pooled libraries were sequenced on an Illumina HiSeq 2500 platform using a 50 bp single read on a high output standard v4 flow cell with a 10–20% PhiX spike-in. About 15 mio reads were obtained for each indexed screen.

For some of the screens, an alternative strategy was used to accommodate sequencing on an Illumina NextSeq 500 platform. Changes to the protocol above include: amplification of extracted gDNA by PCR using a barcoded forward primer (CC_fwd1 to CC_fwd20) and a barcoded reverse primer (CC_Cri_a_rev1 to CC_Cri_a_rev20) using Q5 polymerase (NEB M0491L). Pooled libraries were sequenced on an Illumina NextSeq 500 platform using a 75 bp single read on a high output flow cell with a 2–5% PhiX spike-in. 15-30 mio reads were obtained for each indexed screen.

Sequencing data were analyzed as previously reported^23^ with the following modifications. Trimmed sequences were aligned to the library of protospacers present in the transporter CRISPRi/a sgRNA libraries (Supplementary Table S1), and 89–92% of the number of raw reads typically aligned to the library of protospacers. To estimate technical noise in the screen, simulated negative control genes (the same number as that of real genes) were generated by randomly grouping 10 sgRNAs from the pool of 730 non-targeting control (NTC) sgRNAs present in the libraries. For each gene (and simulated control gene), which is targeted by 10 sgRNAs, two metrics were calculated: (i) the mean of the strongest 7 rho phenotypes by absolute value (“Phenotype score”), and (ii) the p-value of all 10 rho phenotypes compared to the 730 NTC sgRNAs (Mann-Whitney test). To display data compactly, we calculated a single score (“screen score”) by multiplication of the phenotype score with −log10(p-value). sgRNAs were required to have a minimum of 100 counts in at least one of the two conditions tested to be included in the analysis. To deal with the noise associated with potential low count numbers, a pseudocount of 10 was added to all counts. Gene-level phenotype scores and p-values are available in Supplementary Tables S5 and S6.

### Transporter CRISPRi/a screens in rich medium

For K562 CRISPRi/a transporter screens in non-growth limited conditions, pooled libraries were prepared as above. Cells were harvested by centrifugation at 300 ×*g* for 5 min, washed in PBS and resuspended in PBS at 100 mio/mL. Screens were initiated by addition of 7.5 mio cells into 50 mL medium in 150 cm^2^ flasks and were conducted by passaging the libraries in growth medium for 14 days with medium exchange at least each 2 days and keeping the density between 0.1 and 0.4 mio/mL. The pools of cells were harvested at T = 14 days and T = 0 days, and enrichment was analyzed as outlined above. For RPMI + 10% FBS at high confluence, cells were kept at a density of 0.3 – 1.2 mio/mL with medium exchange each 2 days.

For A375 screens, CRISPRi/a parental cells grown in RPMI + 10% FBS in 4× 15 cm cell culture dishes (Falcon 353025) to 80–90% confluence were transduced with transporter library lentiviral supernatant at a MOI of 0.25–0.3 in presence of 8 μg/mL polybrene. Starting 48 hr posttransduction, cells were passaged daily to maintain <95% confluence in RPMI + 10% FBS + puromycin (1.0 μg/mL). After 4 days in puromycin, the proportion of BFP^+^ cells determined by flow cytometry increased to 90–95% of the fraction of viable cells. After recovery for 1 day in puromycin-free medium, library cells were trypsinized and then quenched. Cells were harvested by centrifugation at 300 ×*g* for 5 min, washed in PBS and resuspended in PBS at 100 mio/mL. Screens were initiated by addition of 15 mio cells into 50 mL medium in 15 cm dishes and were conducted by passaging the libraries in growth medium for 14 days with medium exchange at least each 2 days and keeping the confluence between 20–80%. Cells were harvested at T = 14 days and T = 0 days, and enrichment was analyzed as outlined above.

### Transporter screens in subcutaneous tumors in immunodeficient mice

All animal experiments conducted in this study were approved by the MIT Institutional Animal Care and Use Committee (IACUC). A maximum tumor burden of 2 cm was permitted per IACUC protocol, and these limits were not exceeded. Male mice between 3 and 4 months old were used in this study. All animals were housed at ambient temperature and humidity (18–23 °C, 40–60% humidity) with a 12 hr light and 12 hr dark cycle and co-housed with littermates with ad libitum access to water. For animal injections, pooled cells were washed twice with PBS and filtered through a 40 μm cell strainer. After centrifugation at 300 *×g* for 5 min, pelleted cells were resuspended at 100 mio/mL in PBS and the suspensions were kept on ice until injection (<1 hr). NOD.Cg-*Prkdc^scid^ Il2rg^tm1Wjl^*/SzJ mice (NSG mice; The Jackson Laboratory, Strain #005557) were injected subcutaneously into both flanks with 0.1 mL of pooled cell suspension (10 mio cells per tumor). Each screen was conducted with 3 mice (N = 6 tumors) over 14 days. At the endpoint, animals were euthanized and whole tumors (typically 100–500 mg) were excised and kept at −80 °C. gDNA was extracted from homogenized whole tumors (at least 160 μg for each tumor) using the DNeasy Tissue kit (Qiagen). sgRNA barcodes were amplified by PCR using 160 μg of gDNA, as described above for transporter screens in low amino acid screens. Libraries were quantified, pooled and analyzed as described earlier. For each screen, the 2 samples that had the greatest number of NTC counts not within 1 log2 of the median were excluded from the analysis to reduce technical noise. Remaining replicates were paired and read counts were averaged after correcting to match total read counts. These averaged samples were then processed with the T = 0 days samples to determine gene-levels scores as outlined above.

### Determination of engraftment frequency in subcutaneous tumors in NSG mice

Lentivirus was prepared from plasmid pLJM1-eGFP (Addgene #19319). K562 CRISPRi + transporter library cells were infected with pLJM1-eGFP lentivirus and cells expressing GFP were selected by flow cytometry 72 hr post-transduction. GFP^+^ cells were spiked into K562 CRISPRi + transporter library at a ratio of 1:1E3, 1:1E4, or 1:1E5. For each dilution and an unspiked control, cells were prepared as above, diluted with PBS, and 100 μL (0.1, 1 or 10 mio cells) was injected into both flanks of NSG mice. In addition, a portion of each preparation was collected for T = 0 days samples and for growth in RPMI + FBS. Tumors were allowed to form over the course of 19 (10 mio cells injected), 26 (1 mio), and 34 days (0.1 mio). At the endpoint, animals were euthanized and whole tumors were excised and kept on ice. Tumors were dissociated and the presence of GFP^+^ cells was determined by flow cytometry analysis. To determine the skewness of the NTC sgRNA distribution, tumors were analyzed in the same way as the screens.

### Immunoblotting

RPMI media containing low amino acid was made in the same way as for CRISPRi/a screens. In addition, RPMI containing no Arg, His, or Lys was prepared analogously and we used RPMI/rich and RPMI w/o amino acids as controls. For the time course, K562 cells growing in RPMI/rich were harvested, washed twice with treatment medium, and resuspended in treatment medium at a density of 0.35–0.45 mio/mL. After 24 hr, cells were pelleted and resuspended in fresh medium at a density of 0.35–0.45 mio/mL. For each time point, 0.6 mio K562 cells were pelleted by centrifugation at 300 ×*g* for 3 min and flash frozen in liquid nitrogen. Pellets were lysed in 1× RIPA buffer (50 mM Tris HCl, 150 mM NaCl, 0.5% sodium deoxycholate, 0.1% SDS, 1% NP40) and lysates clarified by centrifugation at 21k ×*g* for 10 min at 4 °C. Protein concentration in lysates was quantified by BCA assay (Pierce). 20–25 μg protein was run on a 4–20% Bis-Tris Bolt gel (NuPAGE) in 1× MES buffer (NuPAGE) at 125V for 90 min, and transferred to 0.45 μm nitrocellulose at 18V for 60 min using the Trans-Blot SD semi-dry transfer system (Bio-Rad) in Tris-Glycine transfer buffer (10 mM Tris base, 0.1 M glycine, 20% MeOH). The membrane was blocked with 5% skim milk or BSA (for phospho-antibodies) in TBST for 1 hr at RT, and incubated overnight at 4 °C with primary antibody in the same solution used for blocking (Cell Signaling Technology rabbit antibodies: 1:2000 vinculin (E1E9V) #13901; 1:1000 Phospho-p70 S6 Kinase (Thr389) #9234; 1:1000 p70 S6 Kinase #9202; 1:1000 Phospho-eIF2α (Ser51) (D9G8) #3398; 1:2000 eIF2α #9722; 1:1000 Phospho-4E-BP1 (Ser65) #9451; 1:2000 4E-BP1 (53H11) #9644). After 3× 5 min washes with TBST, incubation for 1–3 hr at RT with 1:5000 secondary antibody (Cell Signaling Technology goat HRP-linked Anti rabbit IgG #7074), and 3 more 5 min washes with TBST, blots were developed in 2 mL Western Lighting Plus chemiluminescent substrate (Perkin Elmer) and imaged on the ImageQuant LAS4000 (Cytiva).

### Growth competition assays

Growth phenotypes were measured in competition assays where a test cell line was mixed with its corresponding NTC control at a 1:1 ratio. Cell lines were grown in complete RPMI and cell densities were determined using a TC20 automated cell counter (Bio-Rad). Cell mixes were prepared by combining 1 mio test cells with 1 mio NTC cells, washing twice with 10 mL PBS, and resuspending the final pellet in 1 mL PBS. 50 μL resuspended cells (0.1 mio) were added to wells of a 12-well plate containing 1 mL of growth medium, and an aliquot was stored at −80 °C to determine the initial ratio for each mix. Cultures were passaged as necessary (i.e daily for low amino acid screen validation) by centrifugation of a portion of the culture and resuspension in fresh medium. At the end of the experiment, the total number of population doublings was determined for all conditions and 0.1–0.2 mio cells were harvested by centrifugation. gDNA was extracted from cell pellets using the QIAamp DNA Blood Mini Kit according to the manufacturer’s instruction, except that the elution was performed using 10 mM Tris·HCl pH 8.5 (typical yields were 2–10 μg). Each extracted gDNA sample was assessed in 2 qPCR reactions using either a forward primer complementary to the test cell sgRNA sequence or to the NTC sgRNA sequence, and a universal reverse primer. Primers were tested beforehand to ensure linearity and specificity of detection. The composition of the mix was estimated by the difference in Ct value between the two reactions (ΔCt). Growth phenotypes were determined by comparing the ΔCt post-growth in the test medium to the ΔCt of the sample taken at T = 0 (ΔΔCt), and then normalizing by the number of population doubling differences between the test medium and a vehicle control (e.g. low amino acid vs RPMI/rich). To determine growth phenotypes in complete media, ΔΔCt values were normalized by the number of population doublings between T = 0 and the end of the assay.

### Amino acid transport assays in K562 cells

The polymer coverslip in a 35 mm μ-Dish (Ibidi 81156) was coated with Cell-Tak. For each dish, 12.25 μg Cell-Tak (Corning 354240) diluted to 360 μL with H_2_O was neutralized with 40 μL 1M bicarbonate pH8 solution and rapidly spread over the surface of the coverslip. After 45 mins at RT, the solution was removed and the coverslip was thoroughly washed with 2× 800 μL H_2_O and subsequently left to dry for a few mins. To prepare cells for immobilization, K562 were harvested (1 mio cells per dish), washed once with RPMI without FBS and resuspended in RPMI without FBS (400 μL per dish). Resuspended cells were added to coated dishes and incubated at RT for 30–45 mins. Medium and unattached cells (typically 0.8 mio cells are needed to form a homogenous monolayer of K562 cells) were gently removed by aspiration. After one wash with 500 μL RPMI/rich, cells were incubated for at least 2 hr at 37 °C/5% CO_2_ in 500 μL RPMI/rich to reach steady state.

For import assays, one dish is required for each time point for each cell line. We typically assayed amino acid import over 7–8 time points (0, 5, 10, 20, 30, 40, 100, 250 s) to capture the initial slope for all 16 amino acids. RPMI/rich was removed by aspiration and was replaced by 400 μL prewarmed and preequilibrated RPMI + 16 heavy-labeled amino acids. After incubation at 37 °C for the duration of the time point (on a heat block for short time points and in the incubator for longer time points), cells were thoroughly washed by sequentially submerging dishes into 4× 400–600 mL ice-cold PBS. For the 0 s time point, the same procedure as above was performed but using RPMI/rich instead of heavy-labeled medium and incubating for 10 s. PBS in the washes was refreshed regularly to ensure limited contamination of intracellular amino acid pools. After the last wash, PBS was removed by aspiration and 600 μL of ice-cold extraction buffer (80% MeOH/20% H_2_O spiked with 4 μg/mL norvaline (Sigma N7627)) was added. Cells were scraped off the plate on ice (United Biosystems MCS–200) and were transferred to 1.5 mL tubes. Samples were homogenized in a thermomixer at 2000 rpm and 4 °C for 15 mins. Tubes were spun at 21.1k ×*g* for 10 min at 4 °C and supernatants were transferred to new tubes containing 10 μL of 0.1N HCl and were stored at −80 °C until analysis by GC-MS. For amino acid consumption assays, a single dish was required for all time points for each cell line. We typically assayed consumption over 5 time points (0, 1, 2, 3, 4 hr) in RPMI and over 4 time points (0, 5, 15, 60 min) in low amino acid medium. RPMI/rich was removed by aspiration and was replaced by 300 μL prewarmed and preequilibrated RPMI/rich. At each time point, 15 μL of medium was removed and kept on ice. All samples were spun at 1000 ×*g* for 5 min at 4 °C and 10 μL of the supernatant was transferred to a tube containing 600 μL of extraction buffer + 10 μL of 0.1N HCl. Samples were stored at −80 °C until analysis by GC-MS.

For GC-MS analysis, samples were dried at RT using a flow of nitrogen. To each tube, we added 24 μL methoxamine (MOX) reagent (ThermoFisher TS–45950) and incubated the resuspended samples at 37 °C for 1 hr. We next added 30 μL N–methyl–N–(tert–butyldimethylsilyl)trifluoroacetamide + 1% tert–Butyldimethylchlorosilane (Sigma 375934) and incubated homogenized samples at 80 °C for 2 hr. After centrifugation at 21.1k ×*g* for 10 min, supernatants were transferred to glass vials. Derivatized samples were analyzed on a DB–35MS column (Agilent Technologies) in an Agilent 7890B gas chromatograph linked to an Agilent 5977B mass spectrometer. 1 μL of sample was injected at 280 °C and mixed with helium carrier gas at a flow rate of 1.2 mL/min. After injection, the oven was held at 100 °C for 1 min and ramped to 250 °C at 3.5 °C/min. The oven was then ramped to 320 °C at 20 °C/min and held for 3 min at 320 °C. Electron impact ionization in the mass spectrometer was performed at 70 eV and the MS source and quadrupole were held at 230 °C and 150 °C, respectively. Scanning mode was used for detection with a scanned ion range of 100–650 m/z. Acetone washes were performed after every 3–5 samples and before and after each run. GC-MS raw data was quantified using El-MAVEN (v0.11.0) and, for each amino acid, we extracted the area of the peak for the fragment with the highest signal over noise for the parent ion and all detectable isotopologues. Natural isotope abundance was subsequently corrected using IsoCorrectoR to determine total ion counts for each amino acid fragment^67^.

For import assays, each sample was first normalized using the norvaline internal standard. We corrected each sample by a factor determined by calculating the ratio of the norvaline signal to the average of the norvaline signal over all time points. As the experiment was run under steady state conditions, we applied a second correction that took advantage of the fact that the total ion count for all amino acid fragments (labeled and unlabeled) remains constant over all time points. We calculated the total ion count (excluding norvaline) for each sample and corrected each sample by a factor determined by calculating the ratio of the total ion count for the sample to the average total ion count over all time points. To determine absolute amounts in each sample, we used a standard mix (Cambridge Isotope Laboratories MSK–A2) containing 17 amino acid to which we added Gln and Asn. We made dilutions of this mix in 0.1N HCl, added them to 600 μL K562 cell extract prepared as for the import assay. We analyzed these samples as above and applied a linear regression to the data. We then used the linear regression data and the mean norvaline signals of the standard and the sample to convert ion counts into pmol amino acid. Data were normalized by the number of cells used in the assay (0.8 mio in cell monolayer minus an estimated 20% cell loss during the washing steps). Intracellular amino acid levels were calculated by averaging the sum of unlabeled and heavy-labeled amino acid over all the time points. To calculate import rates, we determined the initial slope of the increase in intracellular levels of heavy-labeled amino acids. We applied a linear regression to heavy-labeled amino acid levels over time and the number of data points included was variable depending on the amino acid and the conditions. We typically used data from 0–100 s for Asn, Asp, Gln, Glu, Pro; data from 0–40 s for Arg, Gly, His, Ile, Lys, Ser, Thr, Val; data from 0–20 s for Leu, Tyr; and data from 0–10 s for Phe.

For consumption experiments, each sample was normalized using the norvaline standard as above. We next adjusted sample signals to account for the decrease in volume of medium at each time point. Finally, we made use of the fact that the experiment was run under steady state conditions to correct samples based on the total ion count (excluding norvaline) of each sample. We applied a linear regression to the total ion count over the time course of each experiment. We then corrected each sample by a factor calculated by taking the ratio of the total ion count of the sample to the computed level determined by the regression. To determine absolute amounts in each sample, we diluted 10 μL of the standard mix prepared above into 600 μL extraction buffer + 10 μL RPMI/rich and analyzed these standards as detailed for the samples above. We applied a linear regression to the standard data and converted sample ion counts to pmol using the regression data as well as the mean norvaline signals of the standards and samples. Consumption rates were calculated from the slope of the changes in amino acid levels over the time course of the assay determined by linear regression.

For export rates, K562 cells attached in a monolayer on the surface of an Ibidi dish were prepared as for the import assay. Cells were incubated for at least 2 hr with RPMI containing heavy-labeled amino acids at 37 °C/5% CO_2_. Growth medium was removed by aspiration and cells were thoroughly washed by sequentially submerging dishes into 4× 400–600 mL PBS at RT. After aspiration of the last PBS wash, 160 μL prewarmed and preequilibrated RPMI/rich was added to the dish. Plates were kept at 37 °C and the medium was homogenized by regularly swirling the dishes. Samples of the media (12.5 μL) were taken over the course of the assay (0, 5, 10, 20, 30, 40, 100 s) and kept on ice. Samples were centrifuged at 300 ×*g* for 5 min at 4 °C and 10 μL of the supernatant was transferred to a tube containing 600 μL of extraction buffer + 10 μL of 0.1N HCl. Samples were stored at −80 °C until analysis by GC-MS which was performed and analyzed in the same way as the consumption samples.

### Cell viability assays

For K562, cells growing in RPMI/10% FBS were harvested by centrifugation, washed 2× with PBS, and resuspended at 10 mio/mL in PBS. Medium was prepared for each growth condition by addition of small molecules to RPMI low Cys using DMSO stocks. K562 cells were added to each medium to a final concentration of 0.1 mio/mL, and suspensions were dispensed into wells of a 48-well plate in triplicates. After 72 hr incubation at 37 °C/5% CO_2_, 50 μL of each culture was transferred to a 96-well plate (in duplicates), 50 μL CTG reagent was added, and luminescence was measured in a plate reader (BioTek Synergy H1). Viability was quantified by normalization to luminescence at T = 0 hr. For the cystine titration, K562 cells washed in PBS were resuspended in RPMI -Cys (0.1 mio/mL) and dispensed in a 96-well plate. Cystine was added using a D300 digital drug dispenser (HP) and viability at 72 hr was determined by addition of CTG reagent and measurement of luminescence.

For A375, cells were trypsinized and resuspended in RPMI -Cys, then washed 2× with RPMI -Cys, and finally resuspended at 10 mio/mL in the same medium. Cells were added to growth medium to 0.1 mio/mL and dispensed into wells of a 96-well plate. After 2 hr at 37 °C/5% CO_2_, small molecules were added to wells using a D300 digital drug dispenser (HP) using stocks in DMSO. Viability was determined at T = 48 hr by addition of CTG reagent and measurement of luminescence.

For Caco-2 assays, 96-well microplates were coated with poly-D-lysine to avoid cell loss during washing steps. 40 μL of a 1:1 (v/v) solution of poly-D-lysine (Sigma A–003–M) 0.1 mg/mL in H_2_O and PBS was added to each well and incubated for 1 hr at RT. Wells were washed 3× with PBS and left to air dry. Caco-2 cells were seeded into the 96-well plates in EMEM/10% FBS at a density of about 1,000 cells per well. After 48 hr at 37 °C/5% CO_2_, wells were washed twice with RPMI -Cys followed by addition of 50 μL RPMI -Cys. Small molecules were added to wells using a D300 digital drug dispenser (HP). Viability was quantified at T = 48 hr using the CTG luminescence assay and compared to viability at T = 0 hr.

