## Supplementary Figures for "A CRISPRi/a screening platform to study cellular nutrient transport in diverse microenvironments"

#### ADDITIONAL DATA FILES available online

**Supplementary Table S1:** Composition of transporter CRISPRi and CRISPRa sgRNA libraries

**Supplementary Table S2:** Composition of media used in this study

**Supplementary Table S3:** Concentration of amino acid used in low amino acid CRISPRi/a screens and number of population doublings in each screen.

**Supplementary Table S4:** List of oligonucleotides used in this study

**Supplementary Table S5:** CRISPRi screen scores

**Supplementary Table S6:** CRISPRa screen scores

#### SUPPLEMENTARY FIGURES

**Figure S1. Related to Figure 1.** (a) Preparation of K562 CRISPRi or CRISPRa pooled transporter libraries. A custom sgRNA library targeting all annotated transporters (441 solute carriers and 48 ABC transporters) with 10 sgRNAs per gene and 730 non-targeting control (NTC) sgRNAs was constructed and packaged into lentivirus. A K562 monoclonal cell line expressing either the CRISPRi or CRISPRa machinery was infected with lentiviral particles, and untransduced cells were removed by antibiotic selection. Because transporter expression is often tissue-specific[1] and cell lines can lose transporter expression over time[2], CRISPRa bypasses the need for transporter expression in a particular cell line. CRISPRi was used over CRISPR/Cas9 knockout because genetic loss-of-function strategies are often complicated by transcriptional adaptation[3]. For example, SLC7A3 mRNA levels are greatly increased when SLC7A1 is knocked out in mice[4]. In contrast, when the expression of SLC7A1 was strongly reduced using CRISPRi in K562 cells, there was no evidence of compensation by other SLC7 family members (Figure 1B). (b) Assessment of the activity of the mTORC1 and GCN2 pathways in low amino acid screen-like conditions for Arg, Lys, and His via Western blotting of downstream targets eIF2α (GCN2), and S6 kinase and 4E-BP1 (mTORC1). K562 cells grown in complete RPMI medium (Rich; the triangle depicts a centrifugation step) were rapidly pelleted and transferred to either Rich, low amino acid RPMI, or single amino acid dropout RPMI and samples were taken at specific time points after media exchange. For time points longer than 24 hr, the medium was exchanged after 24 hr (medium exchange denoted by asterisk). For all samples, cell lysates were immunoblotted with antibodies against vinculin (loading control), and phospho- and total antibodies against eIF2α, S6K, and 4E-BP1. (c) Tile plots displaying all significant transporter CRISPRi/a hits in low amino acid screens. Screen scores were calculated by multiplying phenotype scores by -log10(p-value) for all genes and computed negative controls. A stringent significance score cutoff was determined by the highest scoring negative control across all conditions for both datasets. All transporter genes that were significant in at least one condition tested are included in the tile plot. Transporter CRISPRi/a cell lines that show a growth defect in RPMI were prone to being pan-resistant in low amino acid conditions, as previously reported in other screens[5], and were removed from the analysis (see Methods). (d) CRISPRi/a transporter screens are highly reproducible as shown by the strong correlation of scores between independent replicates. Black circles represent individual transporter genes and red circles indicate computed negative control pseudogenes assembled from NTC sgRNAs. (e) The custom sgRNA library is highly homogenous, and the library preparation introduces no sample bias. gDNA was isolated from a pool of K562 CRISPRi transporter library cells post-antibiotic selection, and the abundance of each sgRNA present in the library was determined after PCR amplification and high-throughput sequencing (Methods). For this representative sample, >97% of sgRNAs were within 1 log2 of the mean, 12 sgRNAs had less than 1000 counts, and no sgRNA had 0 counts.

**Figure S2. Related to Figure 2.** (a) Cartoon illustrating the competition assay used to validate growth phenotypes of CRISPRi/a cell lines. A CRISPRi/a cell line expressing a test sgRNA is mixed at a 1:1 ratio with cells expressing a non-targeting control (NTC) sgRNA. The ratio of the mix before and after growth in treatment medium is determined by qPCR on gDNA extracted from the culture using a forward primer binding to the test or to the NTC sgRNA and a universal reverse primer. Phenotype scores are determined by normalizing enrichments by the difference in population doublings. Positive values represent cell lines resistant to the slowdown in growth imposed by the treatment medium. (b) Amplification of sgRNAs by qPCR in competition assays is specific. gDNA extracted from K562 CRISPRi/a cells transduced with specific sgRNAs was amplified by qPCR with primers targeting the specific and NTC sgRNA barcodes. Data are the cycle threshold (Ct) value of a representative experiment. Asterisks represent non-specific product amplification. (c) SLC7A8 expression level in K562 SLC7A8 CRISPRi/a cells determined by RT-qPCR relative to the housekeeping gene GAPDH. Error bars: ± SEM. (d) Cartoons illustrating assays performed to measure amino acid import rates and amino acid consumption rates. A monolayer of K562 cells attached to the surface of a Petri dish are incubated in regular medium to reach steady-state. The medium is rapidly exchanged to a similar medium where amino acids are present at the same concentration but are heavy-isotope labeled. After a brief incubation, dishes are thoroughly rinsed with ice-cold PBS and intracellular metabolites are extracted. Light and heavy amino acid levels are quantified using GC-MS. Import rates are determined from the slope of the accumulation of the heavy amino acid over time. The cartoon illustrates the assay using a single amino acid (Arg), but any number of labeled amino acids can be used. Amino acid consumption rates are determined using K562 cells prepared in a similar way. The medium is exchanged to fresh medium and samples are taken over time. Amino acid levels in media samples are determined by GC-MS, and consumption rates are determined by the slope of the change in levels over time. (e) External standard curve used to convert total ion counts determined by GC-MS into absolute levels of amino acids. (f) Representative example of amino acid consumption from the medium of K562 cells growing in RPMI. (g) Data displayed in Figure 2C,D. (h) Comparison of amino acid import rates and inferred export rates in K562 cells growing in RPMI. Import rates are from Figure 2C, and export rates were calculated by subtracting consumption rates from import rates. (i,j) Additional data related to Figure 2F and 2I for amino acids that are not SLC7A5 substrates. (k) Comparison of amino acid import rates in low leucine RPMI and in RPMI for K562 SLC7A5 and NTC CRISPRi determined in Figure 2F. (l) Comparison of intracellular amino acid levels for K562 SLC7A5 and NTC CRISPRi in either low leucine RPMI or in RPMI determined in Figure 2I.

**Figure S3. Related to Figure 3.** (a) Specific changes in intracellular amino acid levels induced by CRISPRi/a of SLC7A1 and SLC7A7. Levels were determined from import assays in Figure 2B (n = 7 ± SEM). (b) Same as a) but for SLC38A3 CRISPRa and data from Figure 2C. (c) CRISPRi/a of genes in the LAT amino acid transporter family leads to specific perturbation of gene expression. LAT1–4 expression level in K562 CRISPRi/a cell lines determined by RT-qPCR relative to the housekeeping gene GAPDH. Error bars: ± SEM of n = 3. (d) Amino acid import rates into K562 cells were influenced by levels of amino acid in the growth medium. Import rates for K562 CRISPRa SLC43A1 and NTC in RPMI and in RPMI modified such that amino acids match human plasma levels (PAA–RPMI) were from Figure 3E. Data represent the mean ratio of import rates in PAA–RPMI compared to RPMI for both cell lines and error bars are the SEM. Amino acid concentrations in the media were determined from their respective formulation. (e) Amino acid import rates into K562 grown in RPMI with one amino acid present at low concentration are specifically reduced only for that amino acid. Data: mean ± SEM of n = 2 (low Val), n = 3 (low Leu), n = 4 (RPMI) of import rates each determined from the linear fit of n ≥ 3 for K562 CRISPRi/a NTC. (f) CRISPRa of SLC43A2 increases import of isoleucine and valine. Amino acid import rates determined in K562 SLC43A2 CRISPRa and SLC43A1 CRISPRi in RPMI (linear fit ± SE of n ≥ 3). Data for SLC43A1 CRISPRa is from Figure 3E. (g) SLC43A2 CRISPRa induces a decrease in intracellular levels of large neutral amino acids in K562 cells in RPMI. Amino acid levels were determined from import assays in (f) (n = 7 ± SEM). (h) SLC43A1 CRISPRa leads to higher export of valine and similar export of leucine, isoleucine, and phenylalanine despite lower intracellular pools. K562 SLC43A1 or NTC CRISPRa cells were grown in RPMI containing heavy-isotope labeled amino acids, rapidly washed with phosphate buffer saline, and incubated in regular RPMI. Accumulation of heavy-labeled amino acids in RPMI was monitored over time by GC-MS.

**Figure S4. Related to Figure 4.** (a) Expression levels of SLC6A4 across cell lines used in this study and specific up- and down-regulation of SLC6A4 via CRISPRi/a. Expression levels of SLC6A4 and SLC7A11 were quantified by RT-qPCR relative to the housekeeping gene GAPDH in K562 CRISPRa, A375 CRISPRa, and Caco-2 CRISPRi cell lines with sgRNAs targeting SLC6A4, SLC7A11 or a non-targeting control (NTC). Error bars: ± SEM of n ≥ 3.

**Figure S5. Related to Figure 5.** (a) CRISPRi transporter screens are highly reproducible as shown by the strong correlation of screen scores between independent replicates. Black circles represent individual transporter genes and red circles indicate computed negative control pseudogenes assembled from NTC sgRNAs. (b) Identification of a larger number of essential transporter genes than in previous screens run in similar conditions. Essential transporters were strongly enriched in expressed genes (98%). Comparison of the number of essential transporter genes in K562 cells growing in RPMI determined in this study to a genome-wide CRISPRi screen[6] (KD: knockdown) and two genome-wide CRISPRko screens all conducted in K562 cells in RPMI[7,8] (KO: knockout). Essential transporter genes were binned by expression level using publicly available data[7]. (c) Pairwise comparison of transporter growth scores determined in this study and in a genome-wide CRISPRi screen[6] in the same cell line, medium, and using the same gRNA library. (d,e) Gene expression levels in K562 CRISPRi cell lines determined by RT-qPCR and relative to the housekeeping gene GAPDH. Error bars: ± SEM of 3 technical replicates. (f) Growth scores of K562 MPC1 and MPC2 CRISPRi determined in 3 growth media and the concentration of a selection of metabolites in those 3 media. Screen scores are from Figure 5C, 6B and metabolites are from published data[9]. (g) Addition of pyruvate to RPMI + FBS or RPMI + dialyzed FBS (dFBS) alleviates the growth defect induced by CRISPRi of MPC1 or MPC2, and addition of lactate to RPMI + dFBS worsens the phenotype. Assays were performed as in Figure 5G and data were normalized to untreated samples. (h) Expression level of SFXN1 in K562 and A375 CRISPRi cell lines determined by RT-qPCR and relative to the housekeeping gene GAPDH. Data: n = 3 ± SEM.

**Figure S6. Related to Figure 6.** (a) Determination of the engraftment frequency of K562 cells injected subcutaneously in immunodeficient mice. K562 CRISPRi library cells expressing GFP were mixed to pools of K562 CRISPRi cells at a ratio ranging from 1:1000–1:100,000. 10, 1 or 0.1 mio cells of these preparations were injected into the flank of NSG mice. Tumors formed over 19–34 days, were harvested and dissociated into single cell suspensions. The presence (or absence) of $GFP^+$ cells in homogenized tumors was detected by flow cytometry and was used to determine the engraftment frequency and optimize conditions to preserve library complexity during in vivo transporter screens. (b) Analysis of the distribution of counts for all non-targeting control (NTC) sgRNA is used to assess the quality of transporter screens. Plot displaying the number of counts for all NTC sgRNAs determined from the sequencing of tumor samples from (a), the initial pool of library cells, and library cells grown in RPMI. (c) Expression level of SLC2A1 in K562 and A375 CRISPRa cell lines determined by RT-qPCR and relative to the housekeeping gene GAPDH. Data: n = 3 ± SEM.

### FIGURE S1

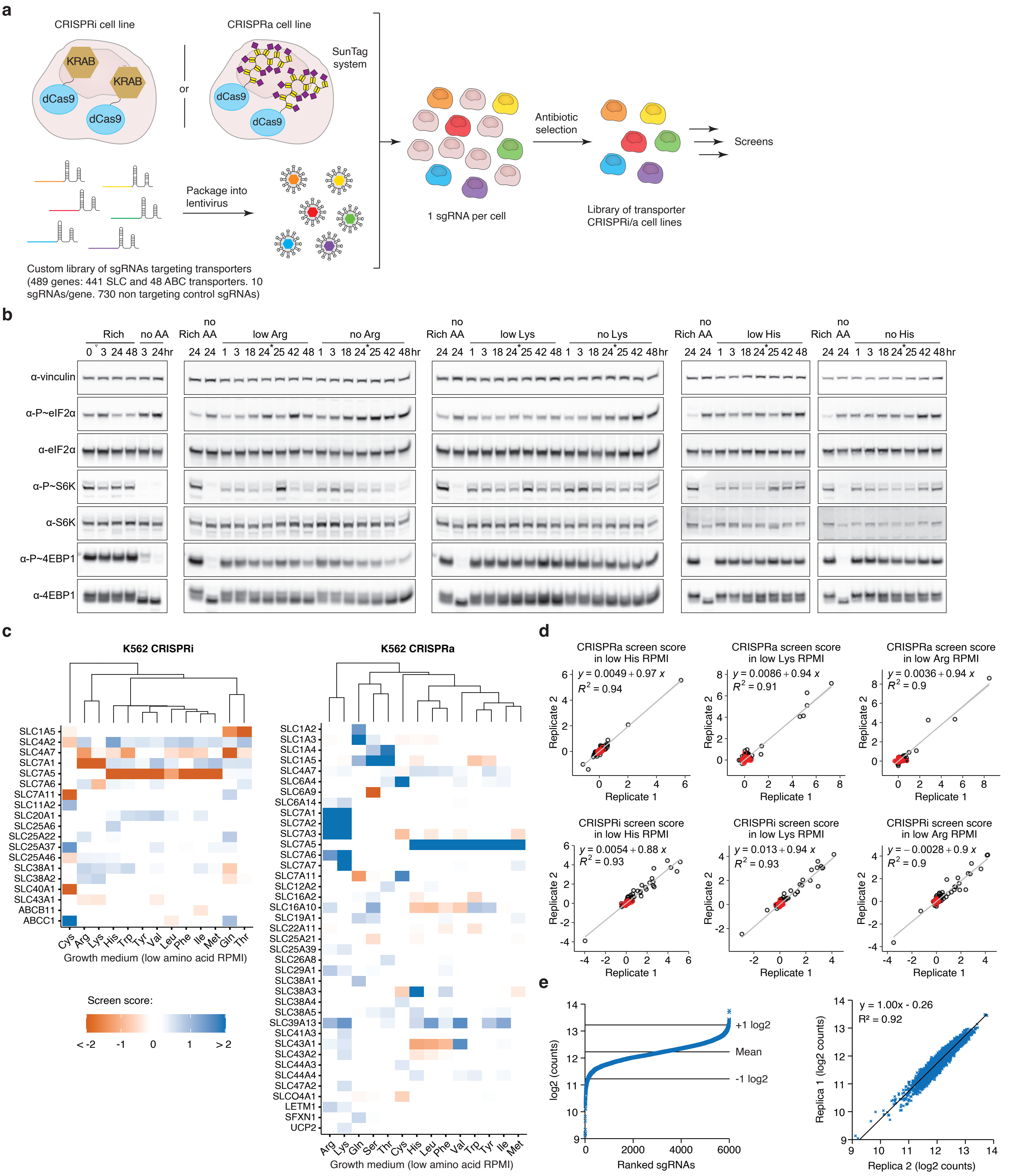
a
CRISPRi cell line
CRISPRa cell line
KRAB
dCas9
or
SunTag system
Custom library of sgRNAs targeting transporters (489 genes: 441 SLC and 48 ABC transporters. 10 sgRNAs/gene. 730 non targeting control sgRNAs)
Package into lentivirus
1 sgRNA per cell
Antibiotic selection
Library of transporter CRISPRi/a cell lines
Screens
b
Rich
no AA
no Rich AA
low Arg
no Arg
low Lys
no Lys
low His
no His
α-vinculin
α-P~eIF2α
α-eIF2α
α-P~S6K
α-S6K
α-P~4EBP1
α-4EBP1
c
K562 CRISPRi
K562 CRISPRa
Growth medium (low amino acid RPMI)
Screen score:
< -2
-1
0
1
> 2
d
CRISPRa screen score in low His RPMI
$y = 0.0049 + 0.97\ x$
$R^2 = 0.94$
CRISPRa screen score in low Lys RPMI
$y = 0.0086 + 0.94\ x$
$R^2 = 0.91$
CRISPRa screen score in low Arg RPMI
$y = 0.0036 + 0.94\ x$
$R^2 = 0.9$
CRISPRi screen score in low His RPMI
$y = 0.0054 + 0.88\ x$
$R^2 = 0.93$
CRISPRi screen score in low Lys RPMI
$y = 0.013 + 0.94\ x$
$R^2 = 0.93$
CRISPRi screen score in low Arg RPMI
$y = -0.0028 + 0.9\ x$
$R^2 = 0.9$
Replicate 1
Replicate 2
e
log2 (counts)
Ranked sgRNAs
+1 log2
Mean
-1 log2
y = 1.00x - 0.26
R² = 0.92
Replica 1 (log2 counts)
Replica 2 (log2 counts)

#### FIGURE S2

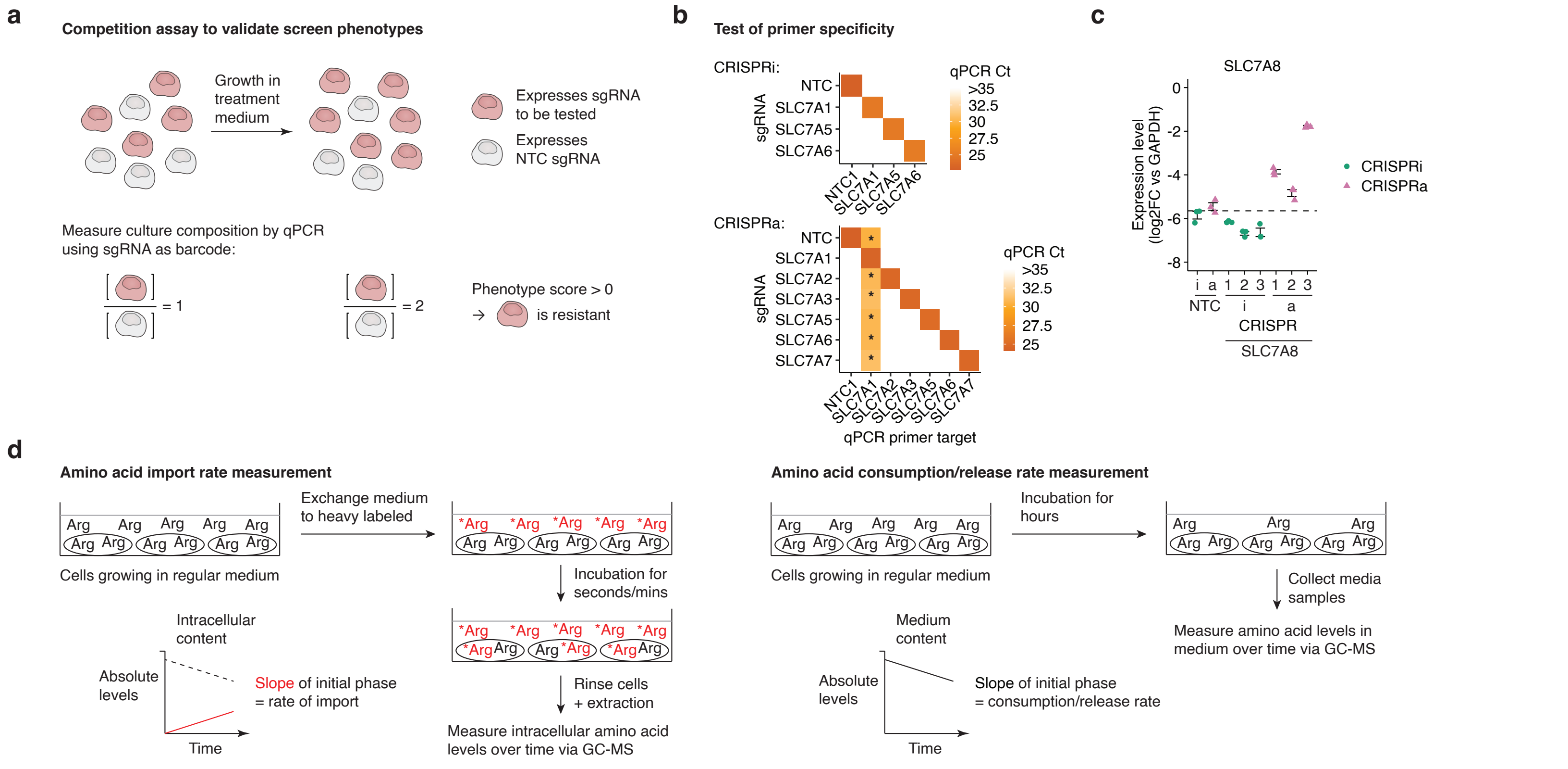


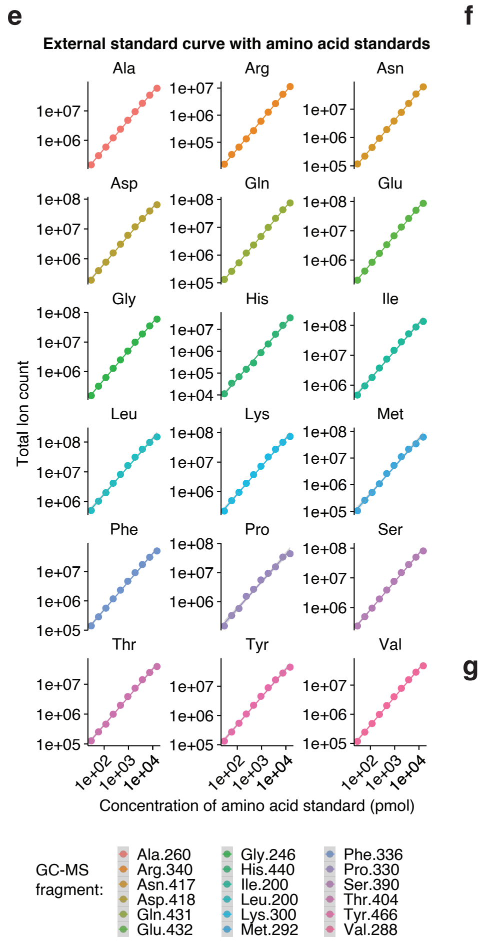


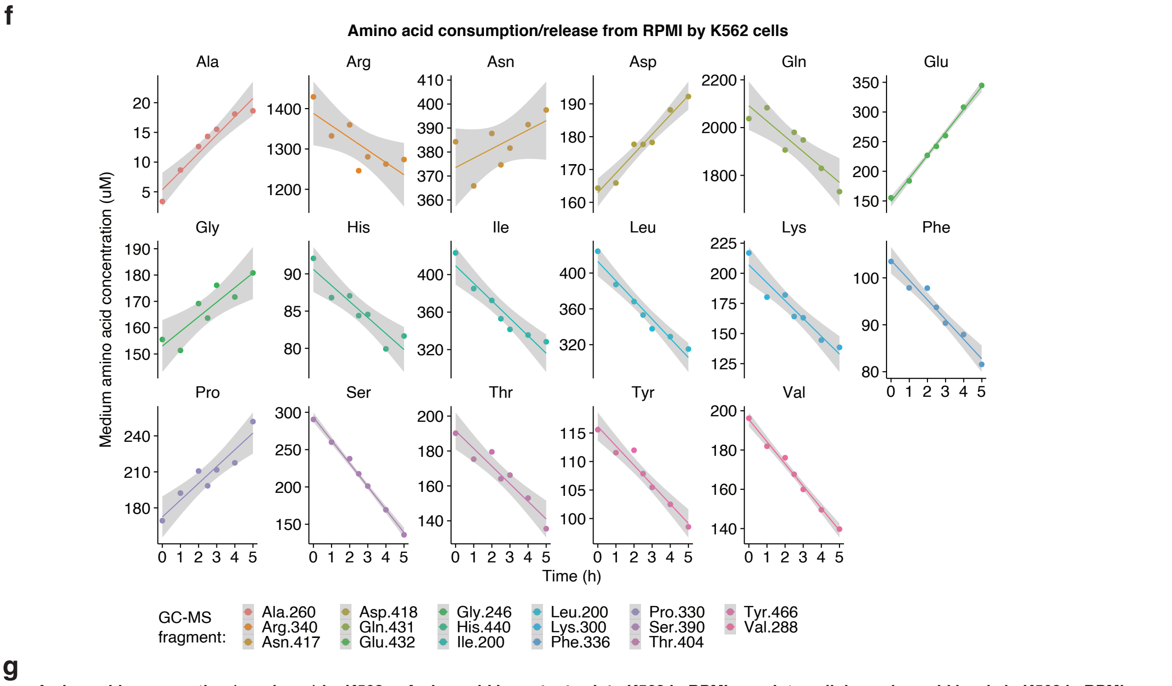


**g**

**Amino acid consumption (or release) by K562 in RPMI** (fmol/hr per cell; ave of 5 experiments)

| | | | |
|---|---|---|---|
| Arg | Asn | Asp | Gln |
| -1 +/- 3 | 2.2 +/- 0.3 | 2.0 +/- 0.3 | -25 +/- 2 |
| Glu | Gly | His | Ile |
| 14 +/- 1 | 3.5 +/- 0.7 | -0.6 +/- 0.1 | -9 +/- 1 |
| Leu | Lys | Phe | Pro |
| -10 +/- 1 | -4.1 +/- 0.4 | -1.9 +/- 0.2 | 5 +/- 2 |
| Ser | Thr | Tyr | Val |
| -13 +/- 2 | -5.0 +/- 0.4 | -1.3 +/- 0.1 | -5 +/- 1 |

**Amino acid import rates into K562 in RPMI** (fmol/hr per cell; average of 6 experiments)

| | | | |
|---|---|---|---|
| Arg | Asn | Asp | Gln |
| 52 +/- 9 | 52 +/- 10 | 0.3 +/- 0.2 | 460 +/- 110 |
| Glu | Gly | His | Ile |
| 33 +/- 3 | 4.0 +/- 0.9 | 16 +/- 2 | 65 +/- 8 |
| Leu | Lys | Phe | Pro |
| 172 +/- 22 | 4.0 +/- 0.8 | 74 +/- 6 | 2.8 +/- 0.5 |
| Ser | Thr | Tyr | Val |
| 90 +/- 7 | 54 +/- 4 | 39 +/- 2 | 10.9 +/- 1.3 |

**Intracellular amino acid levels in K562 in RPMI** (fmol per cell; average of 7 experiments)

| | | | |
|---|---|---|---|
| Arg | Asn | Asp | Gln |
| 1.3 +/- 0.1 | 7.2 +/- 0.9 | 9.6 +/- 0.7 | 30 +/- 6 |
| Glu | Gly | His | Ile |
| 36 +/- 1 | 8.4 +/- 1.4 | 0.5 +/- 0.1 | 1.7 +/- 0.1 |
| Leu | Lys | Phe | Pro |
| 1.9 +/- 0.2 | 0.20 +/- 0.04 | 0.46 +/- 0.04 | 6.1 +/- 0.8 |
| Ser | Thr | Tyr | Val |
| 3.8 +/- 0.3 | 2.7 +/- 0.2 | 0.66 +/- 0.07 | 0.64 +/- 0.05 |
| Ala | | | |
| 0.14 +/- 0.02 | | | |

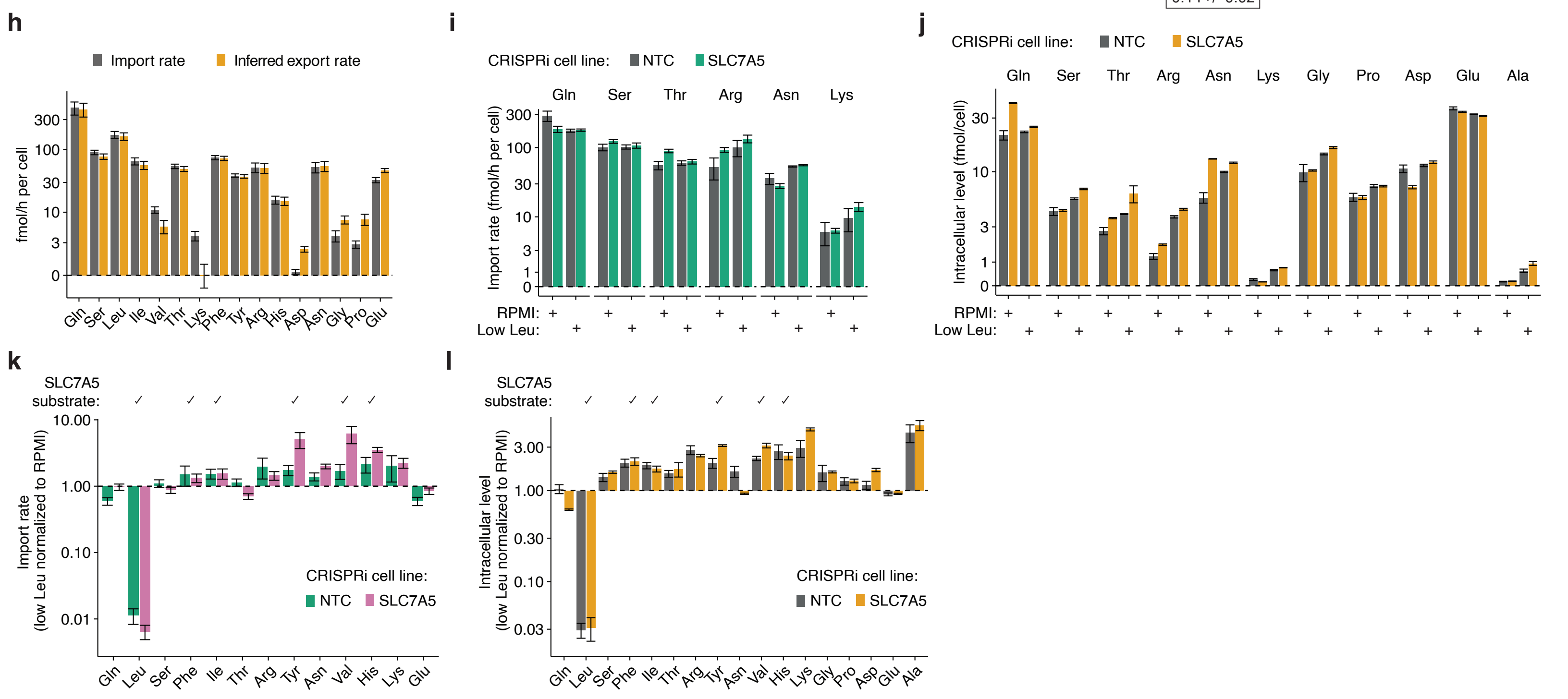

FIGURE S3

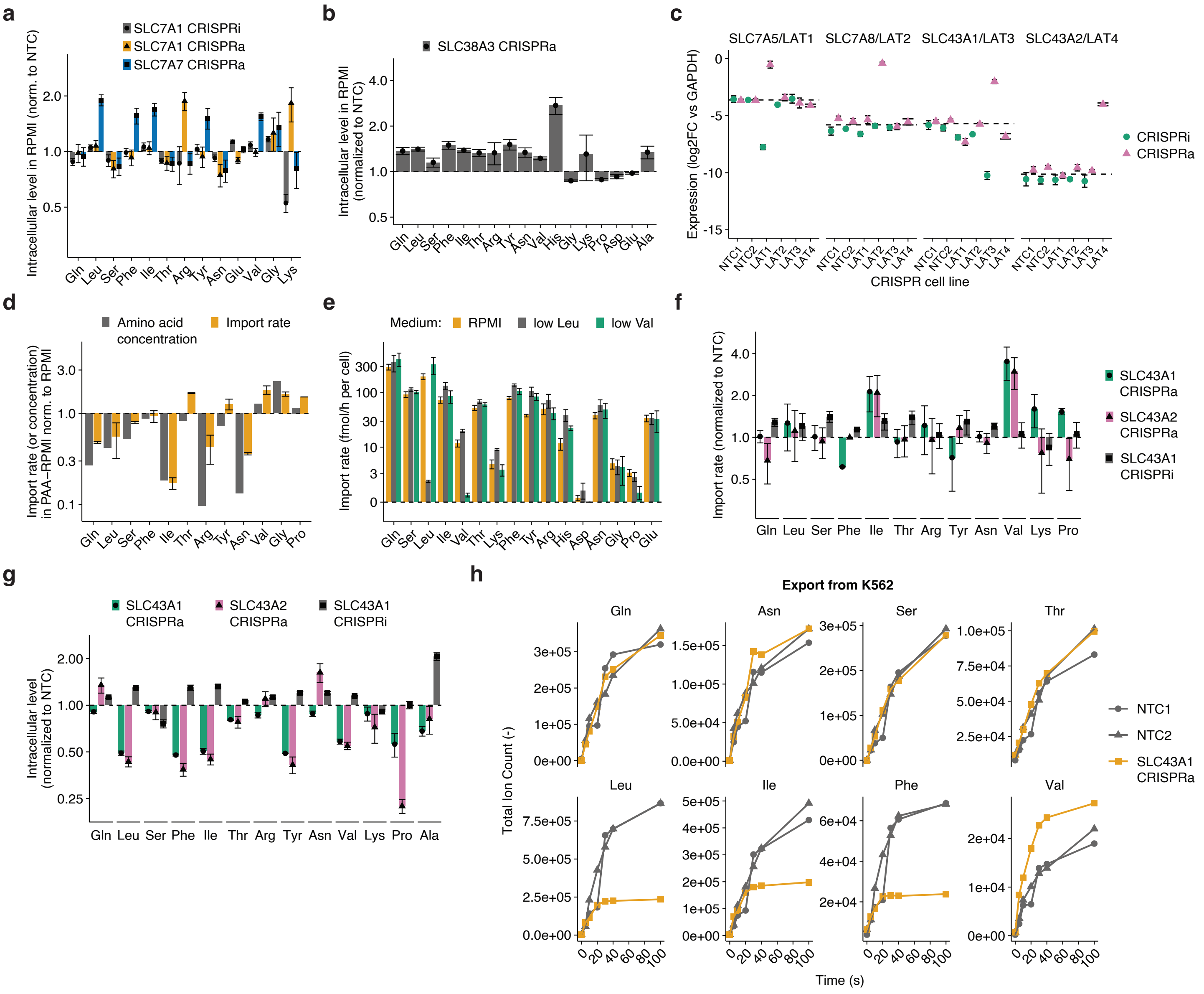
a
SLC7A1 CRISPRi
SLC7A1 CRISPRa
SLC7A7 CRISPRa
Intracellular level in RPMI (norm. to NTC)
b
SLC38A3 CRISPRa
Intracellular level in RPMI (normalized to NTC)
c
SLC7A5/LAT1
SLC7A8/LAT2
SLC43A1/LAT3
SLC43A2/LAT4
Expression (log2FC vs GAPDH)
CRISPRi
CRISPRa
CRISPR cell line
d
Amino acid concentration
Import rate
Import rate (or concentration) in PAA-RPMI norm. to RPMI
e
Medium: RPMI low Leu low Val
Import rate (fmol/h per cell)
f
SLC43A1 CRISPRa
SLC43A2 CRISPRa
SLC43A1 CRISPRi
Import rate (normalized to NTC)
g
SLC43A1 CRISPRa
SLC43A2 CRISPRa
SLC43A1 CRISPRi
Intracellular level (normalized to NTC)
h
Export from K562
Gln
Asn
Ser
Thr
Leu
Ile
Phe
Val
Total Ion Count (-)
Time (s)
NTC1
NTC2
SLC43A1 CRISPRa

#### FIGURE S4

**a**

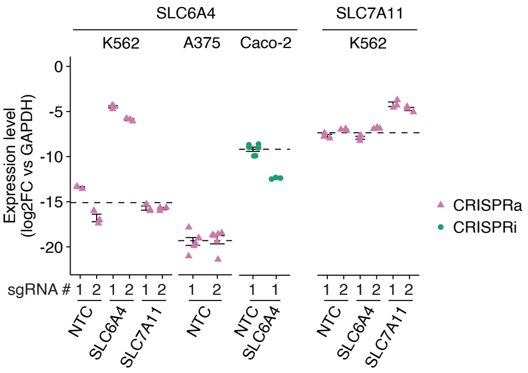

SLC6A4
SLC7A11
K562
A375
Caco-2
K562
Expression level (log2FC vs GAPDH)
0
-5
-10
-15
-20
CRISPRa
CRISPRi
sgRNA #
1 2 1 2 1 2
1 2
1 1
1 2 1 2 1 2
NTC
SLC6A4
SLC7A11
NTC
NTC
SLC6A4
NTC
SLC6A4
SLC7A11

### FIGURE S5

a

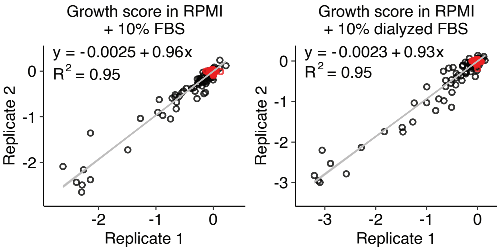


b

K562 essential genes in RPMI + 10% FBS

| | | This study (CRISPR KD) | Horlbeck et al. (CRISPR KD) | To et al. (CRISPR KO) | Wang et al. (CRISPR KO) | Total # of genes |
|---|---|---|---|---|---|---|
| Transporter genes | | 46 | 29* | n.a. | 20* | 489 |
| Expression | High | 42 | 27 | 29* | 17 | 258 |
| | Low | 3 | 2 | n.a. | 1 | 130 |
| | N.d. | 1 | 0 | n.a. | 2 | 101 |
| *Overlap with this study: | | | 24 | 19 | 13 | |

c

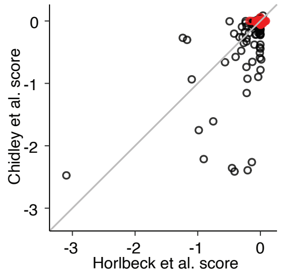


d

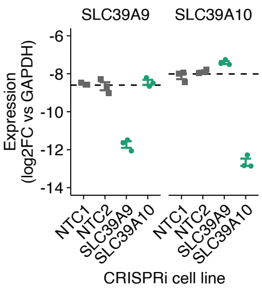


e

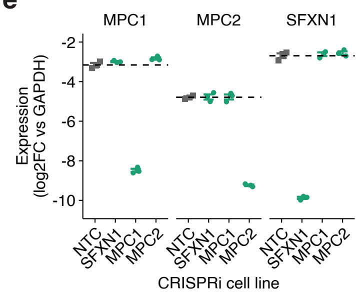


f

| CRISPRi screen scores | | RPMI + 10% FBS | RPMI + 10% dFBS | HPLM + 10% dFBS |
|---|---|---|---|---|
| | MPC1 | -0.03 | -1.39 | -0.01 |
| | MPC2 | -0.08 | -1.61 | 0.00 |

| Metabolite concentration in medium (µM) | | RPMI + 10% FBS | RPMI + 10% dFBS | HPLM + 10% dFBS |
|---|---|---|---|---|
| | Pyruvate | 2.7 | N/D | 42.6 |
| | Lactate | 1512 | 10.7 | 1524 |
| | Alanine | 134 | 2.7 | 617 |
| | Glucose | 5241 | 4929 | 5110 |
| | Glutamate | 172 | 101 | 73 |

g

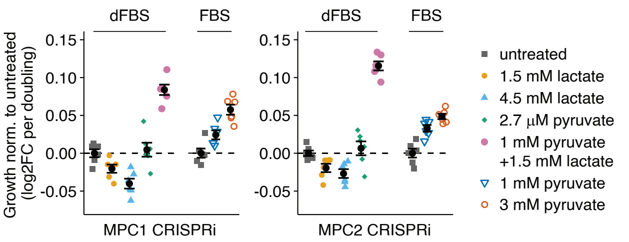


h

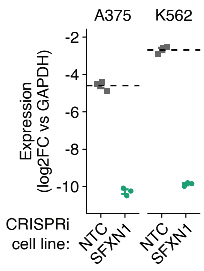

### FIGURE S6

**a**

**Determination of engraftment frequency in K562:**

Subcutaneous injection into recipient mice (2 flanks per condition)

in vivo growth

Dissociation of tumor cells and detection of GFP⁺ cells by flow cytometry

**Proportion of GFP⁺ cells:**

| **# of cells injected:** | 1:1000 GFP⁺ cells | 1:10,000 GFP⁺ cells | 1:100,000 GFP⁺ cells | 0 GFP⁺ cells |
|---|---|---|---|---|
| 10 mio | **10k GFP⁺ cells** | **1k GFP⁺ cells** | **100 GFP⁺ cells** | 0 GFP⁺ cells |
| 1 mio | **1k GFP⁺ cells** | **100 GFP⁺ cells** | 10 GFP⁺ cells | 0 GFP⁺ cells |
| 0.1 mio | 100 GFP⁺ cells | 10 GFP⁺ cells | 1 GFP⁺ cell | 0 GFP⁺ cells |

**Detection of GFP⁺ cells in tumor samples?**

| | | | |
|---|---|---|---|
| **YES** | **YES** | **YES** | No |
| **YES** | **YES** | No | No |
| No | No | No | No |

→ **Engraftment rate of >1% when 1 or 10 mio cells are injected (<1% when 100k cells are injected)**

**b**

**Distribution of non-targeting control (NTC) sgRNAs in samples**

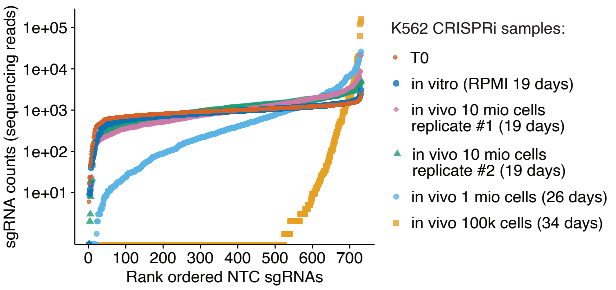


**c**

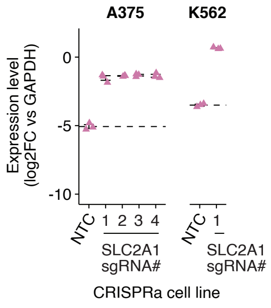
